# Restoring the loss of consciousness under sevoflurane anesthesia in states of anxiety and depression by potently activating serotonergic pathways in brain

**DOI:** 10.1101/2024.06.19.599720

**Authors:** YuLing Wang, Yue Yang, YaXuan Wu, XiaoXia Xu, WeiHui Shao, Shuai Li, Qing Xu, LeYuan Gu, Lu Liu, ZiWen Zhang, XuanYi Di, Hui Wu, DanDan Yan, Yue Shen, XinZhong Chen, HongHai Zhang

**Affiliations:** Department of Anesthesiology, Affiliated Hangzhou First People’s Hospital, Westlake University School of Medicine, Hangzhou, 310006, China; Department of Anesthesiology, the Fourth Clinical School of Medicine, Zhejiang Chinese Medical University, Hangzhou, 310006, China; Department of Anesthesiology, National Cancer Center/National Clinical Research Center for Cancer/Cancer Hospital, Chinese Academy of Medical Sciences and Peking Union Medical College, Beijing 100021, China; Department of Anesthesiology, Zhejiang University School of Medicine, Hangzhou, 310006, China; Department of Anesthesia, Women’s Hospital, Zhejiang University School of Medicine, Hangzhou, 310006, China; Westlake Laboratory of Life Sciences and Biomedicine, Hangzhou, 310006, China

**Author notes:** These authors contributed equally to this work (YuLing Wang, Yue Yang, YaXuan Wu, XiaoXia Xu, WeiHui Shao, Shuai Li). Corresponding authors: XinZhong Chen, HongHai Zhang.

**Keywords:** General anesthesia, Delayed emergence, Sevoflurane, 5-hydroxytryptamine, Central amygdala nucleus, Medial prefrontal cortex, Consciousness, Anxiety, Depression

## Abstract

Preoperative anxiety and depression are significant risk factors for prolonged general anesthesia emergence, raising postoperative complications and mortality. However, the neural mechanisms underlying this association remain unclear, limiting effective interventions and development of effective preventive and therapeutic strategies. Previous studies have shown that sleep-wakefulness neural circuit is closely related to neural circuitry of anesthesia recovery, with 5-Hydroxytryptamine (5-HT) playing a pivotal role of anesthesia arousal. Herein, we utilized pharmacology, optogenetics, chemogenetics, fiber photometry and retrograde tracing to demonstrate that activation of the 5-HTergic neural circuit between the central amygdala (CeA) and medial prefrontal cortex (mPFC) accelerates recovery from sevoflurane anesthesia. Particularly for individuals with anxiety- and depression-like symptoms, who exhibit more severe delayed emergence, it is necessary to enhance 5-HT neuronal activity by further optimizing parameters. To further validate the reliability of our findings and the specificity of viral tools, we successfully bred TPH2-Cre transgenic mice and repeated key experiments, which robustly confirmed all original conclusions. Pharmacological experiments further demonstrated that both 5-HT₁A and 5-HT₂A receptors mediate the pro-arousal function of this circuit. Collectively, our study identifies a key CeA-mPFC serotonergic pathway linking emotional disorders to abnormal anesthetic emergence, providing potential targets for optimizing anesthesia management in vulnerable patients.

**Highlights:**

- Boosting the 5-HTergic neural circuit between the CeA and mPFC promotes emergence from sevoflurane anesthesia, with 5-HT_1_A and 5-HT_2_A receptors playing a critical role in modulating consciousness transitions.
- Preoperative anxiety and depression exacerbate delayed emergence from anesthesia, highlighting the need for targeted interventions.
- Deciphering the aberrant neurocircuitry mechanisms underlying delayed emergence to inversely deduce the neurocircuitry principles of normal physiology represents a pathological-to-physiological research paradigm.
- Stepwise enhancement of intervention parameters reverses delayed emergence in pathological states.
- Tailored activation of the 5-HTergic circuit offers a novel approach to mitigate complications from delayed emergence in vulnerable patient populations.

**Graphical Abstract:** 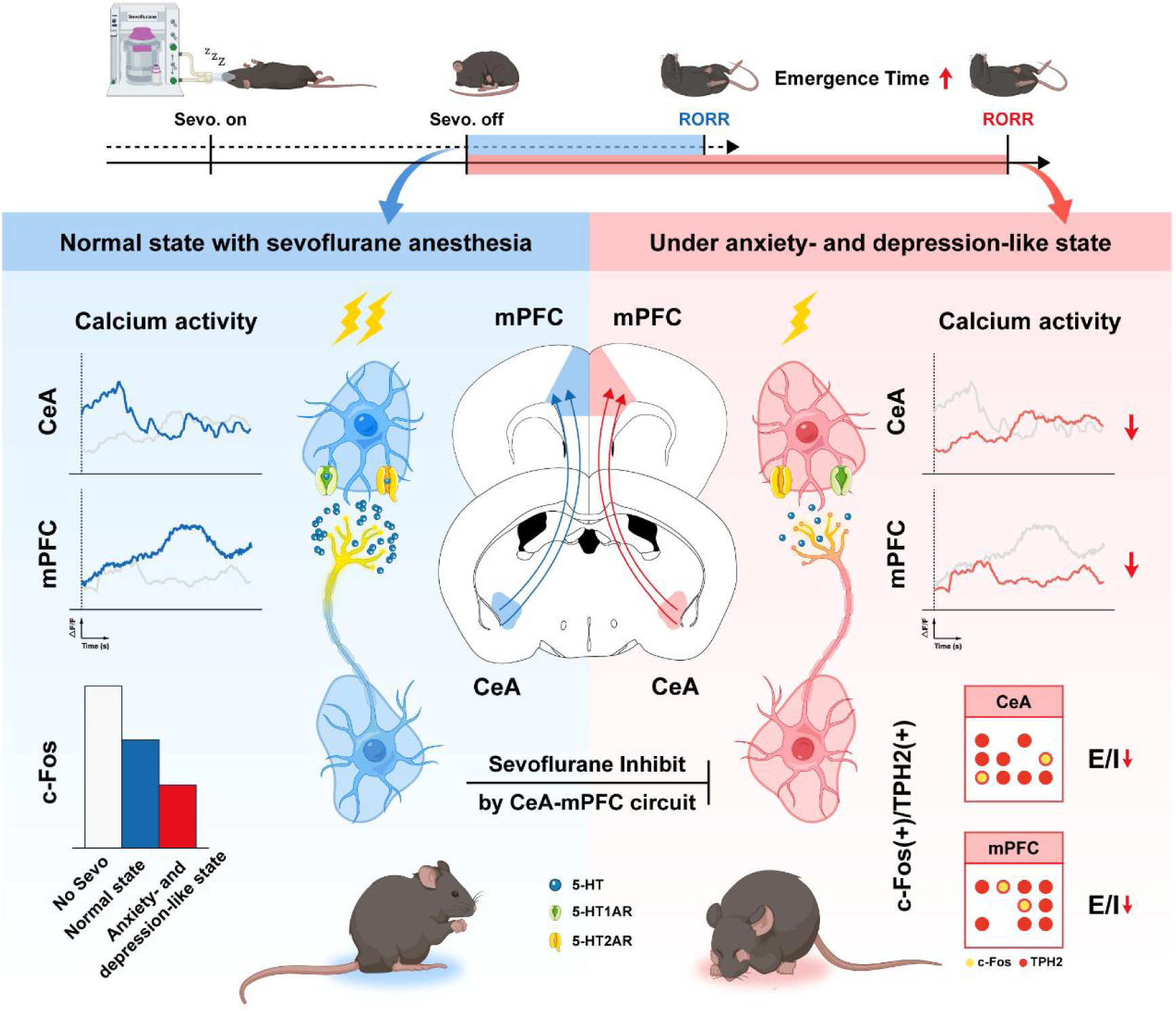

## 1 Introduction

Anxiety and depression represent significant global public health burdens, with prevalence rising by 26% and 28%, respectively, following the COVID-19 pandemic^1^. These conditions are highly prevalent among surgical patients and are clinically significant predictors of adverse perioperative outcomes^2^. Preoperative anxiety and depression are correlated with more severe delayed emergence from anesthesia, a phenomenon linked to alterations in the neural circuits regulating the sleep-wake cycle regulation^3,4^. Given these risks, understanding the mechanisms linking psychological factors to anesthesia recovery is crucial for improving perioperative outcomes. Only through a meticulous exploration of these factors and the implementation of targeted interventions can we hope to reduce the risks associated with delayed emergence from anesthesia, thereby enhancing overall patient safety and reducing the incidence of adverse medical events^5,6^.

General anesthesia represents a meticulously adjusted, drug-induced, controllable state that guarantees patient comfort and safety during surgical and invasive clinical procedures^7^. Among the myriad anesthetics available, sevoflurane, a potent inhalational anesthetic, has emerged as a cornerstone in modern surgery due to its rapid induction and smooth emergence^8^. However, despite its widespread use, the exact mechanisms by which sevoflurane affects the brain, particularly its intricate role in facilitating arousal from anesthesia, remain shrouded in a degree of mystery^9,10^. Notably, delayed emergence has also become a common complication of general anesthesia^11^. Yet, failure to regain consciousness promptly after anesthesia can lead to severe consequences, such as increased risks of hypoxemia, respiratory suppression, metabolic encephalopathy, and central nervous system damage, with potentially life-threatening outcomes^12–15^. The causes of delayed emergence from general anesthesia are complex. Studies have shown that preoperative psychological status can be a potential predictor of the need for anesthesia. Preoperative anxiety and depression, two predominant psychological comorbidities in surgical patients, are closely correlated with prolonged emergence duration and elevated risks of postoperative adverse complications^16,17^.

5-Hydroxytryptamine (5-HT), commonly referred to as serotonin, is a crucial monoamine neurotransmitter ubiquitously present in mammalian tissues^18,19^. Its role spans numerous aspects of central nervous system function, including cognitive processing, emotional regulation, sexual activity, thermoregulation, and even the regulation of food consumption. Studies have demonstrated that central 5-HT levels fluctuate dynamically during anesthesia, hinting at its pivotal role in modulating the effects of general anesthesia^20,21^. In particular, patients with anxiety-depressive disorders usually have dysfunctional 5-HTergic neurons, resulting in a reduced regulation of their neural networks. Such neuronal impairment impairs neural network modulation, blunts responsiveness to arousal cues, and ultimately alters anesthetic emergence profiles. In addition, 5-HT imbalance is closely associated with dysfunction of the autonomic nervous system, which may exacerbate autonomic instability during postoperative recovery, leading to additional complications such as cardiovascular instability and respiratory depression^22–24^. The cerebral cortex and the dorsal raphe nucleus (DR) boast a significant concentration of 5-HT neurons. Serotonergic neurons originating from DR project extensively to the midbrain and forebrain, participating in the composition of the central ascending arousal system^25,26^. Meanwhile, the cerebral cortex, limbic system, basal forebrain, amygdala, and other nuclei rich in cholinergic and monoaminergic neurons also project back to the DR, creating a sophisticated network that orchestrates the intricate balance between sleep and wakefulness^27–29^. The central amygdala nucleus (CeA) plays a crucial role in various behaviors, including those related to fear, anxiety, and depression^30,31^. It receives substantial inputs from the cerebral cortex and subcortical areas, and in turn, projects to key brainstem and diencephalic structures, as well as to regions engaged in hormonal, autonomic, and stress-related responses, thereby influencing the sleep-wake cycle^31^. Additionally, CeA also sends numerous projections to areas involved in arousal and rapid eye movement sleep, such as the pons-geniculate-occipital waves, further emphasizing its indispensable role in regulating sleep and wakefulness^32,33^.

The medial prefrontal cortex (mPFC) is intricately connected with arousal-related nuclei, including the locus coeruleus (LC) and DR^11,34^. This structural and functional entanglement indicates that the mPFC is pivotal in modulating consciousness levels, not only during anesthesia but also in the transition from anesthesia to wakefulness^35–37^. Recent research has demonstrated that inhibiting mPFC neurons prolongs the emergence of consciousness from general anesthesia, further solidifying its role as a key regulator of arousal and consciousness^38^. Additionally, studies have observed a decrease in serotonin levels in the frontal cortex during anesthesia, which may contribute to alterations in consciousness associated with anesthetic states. Researches have revealed widespread neural projections between the CeA and the mPFC, yet the role of the 5-HT neural pathway within the CeA-mPFC circuit during the process of emergence from general anesthesia remains unclear^39,40^. Therefore, this study aims to explore the involvement of 5-HT neurons in the CeA and mPFC, as well as their functional connectivity, in the restoration of consciousness during sevoflurane anesthesia. Particularly, the functional activity of this neural circuit under different degrees of delayed emergence is both intriguing and significant.

In this study, we investigated the impact of CeA^5-HT^ and mPFC^5-HT^ neurons on sevoflurane anesthesia emergence time through exogenous 5-hydroxytryptophan (5-HTP) supplementation and endogenous optogenetic and chemogenetic modulation. Using multidisciplinary approaches, we delineate the neurocircuitry and physiological functions between CeA and mPFC. To further validate the reliability of our core conclusions and the specificity of viral tools used in this work, we successfully bred TPH2-Cre transgenic mice and repeated key functional experiments, which solidly verified all our major findings. Our findings demonstrate that by activating the CeA-mPFC serotonergic pathway, consciousness can be effectively restored from sevoflurane anesthesia, especially in cases of anxiety and depression, providing a targeted neurotherapeutic strategy for this high-risk population. Consequently, this article contributes to the understanding of the underlying neural circuit activity involved in anesthesia emergence and provides potential intervention strategies to improve postoperative outcomes in clinical patients, especially those with preoperative anxiety and depression, by mitigating delayed emergence and preventing adverse complications.

We further clarify the translational relevance of the animal models adopted in this study: (1) The chronic restraint stress (CRS) model simulates chronic stress and is consistent with the neuropathological mechanisms of clinical anxiety disorders (such as hypothalamic-pituitary-adrenal axis activation); (2) The lipopolysaccharide (LPS)-induced model simulates inflammation-related depression, which is associated with the common postoperative inflammatory state in perioperative patients; (3) Although these two models cannot fully replicate human diseases, they can cover the key pathological features of clinical anxiety and depression, providing a reasonable animal model basis for this study.

## 2 Results

### 2.1 Anxiety- and depression-like states prolong sevoflurane anesthesia emergence

To clarify how preoperative affective disorders alter consciousness recovery after sevoflurane anesthesia, we established two classic mouse pathological models: chronic restraint stress (CRS)-induced and lipopolysaccharide (LPS)-induced anxiety/depression models. A full battery of behavioral paradigms including open field test (OFT), elevated plus maze (EPM), forced swim test (FST) and tail suspension test (TST validated robust anxiety- and depressive-like behaviors in both CRS and LPS mice (Extended Data Fig. 1A-H). Compared with naive control animals, CRS mice exhibited markedly reduced central zone exploration and open-arm entries, alongside prolonged immobility in despair tests (P < 0.001, P < 0.01), confirming successful model construction. Subsequent sevoflurane anesthesia assays demonstrated unaltered induction latency but significantly prolonged emergence time in both CRS and LPS groups relative to wild-type controls (P < 0.01; Fig. 1A-B). These data indicate that pathological anxiety and depression selectively disrupt arousal processes without affecting anesthetic onset. Accordingly, we systematically dissected the underlying serotonergic circuit remodeling driving delayed emergence under emotional dysfunction.

**Figure 1.**
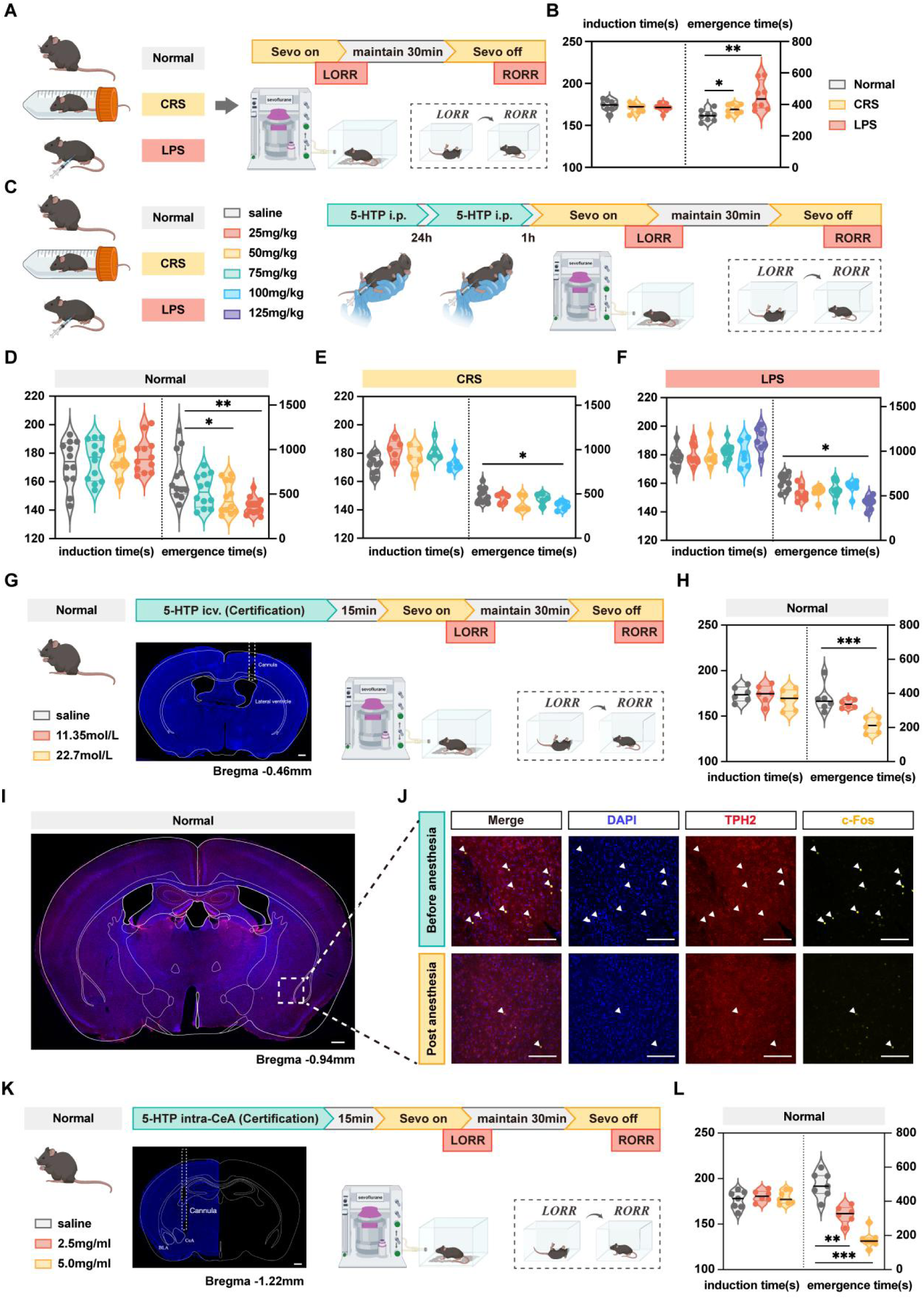
Anxiety- and depression-like states delay emergence from sevoflurane anesthesia, while exogenous supplementation of 5-HTP facilitates emergence. **(A)** Experimental protocol for sevoflurane anesthesia and monitoring the duration of induction and emergence time. **(B)** Induction time and emergence time after sevoflurane anesthesia for the Normal, CRS and LPS groups. **(C)** Experimental protocol for i.p. injection of different doses of 5-HTP and monitoring the duration of induction and emergence time. **(D)** Induction time and emergence time for normal mice for i.p. injection in the saline control, 25 mg/kg 5-HTP, 50 mg/kg 5-HTP, and 75 mg/kg 5-HTP groups. **(E)** Induction time and emergence time for CRS mice for i.p. injection in the saline control, 25 mg/kg 5-HTP, 50 mg/kg 5-HTP, 75 mg/kg 5-HTP and 100 mg/kg 5-HTP groups. **(F)** Induction time and emergence time for LPS mice for i.p. injection in the saline control, 25 mg/kg 5-HTP, 50 mg/kg 5-HTP, 75 mg/kg 5-HTP, 100 mg/kg 5-HTP and 125 mg/kg 5-HTP groups. **(G)** Experimental protocol for icv. injection of different doses of 5-HTP, and representative image of the track of cannula implanted into the lateral ventricle. **(H)** Induction time and emergence time for normal mice for icv. injection in the saline control, 11.35 mol/L 5-HTP, and 22.70 mol/L 5-HTP groups. **(I)** Representative coronal brain slice of CeA, showing the staining for c-Fos, TPH2 and DAPI before anesthesia. **(J)** Representative images of staining for c-Fos, TPH2, and DAPI in the CeA before or after anesthesia. **(K)** Experimental protocol for intra-CeA microinjection of 5-HTP, and representative image of the track of cannula implanted into the CeA. **(L)** Induction time and emergence time for intra-CeA microinjection in the saline control, 2.5 mg/ml 5-HTP, and 5.0 mg/ml 5-HTP groups. ***p < 0.001; **p < 0.01; *p < 0.05; Scale bar: 100 μm; Data are mean ± SEM; i.p., intraperitoneal injection; icv., intracerebroventricular injection; 5-HTP, 5-hydroxytryptophan; CeA, central amygdala nucleus; Sevo, sevoflurane; LORR, loss of righting reflex; RORR, recovery of righting reflex; CRS, chronic restraint stress; LPS, lipopolysaccharide

### 2.2 Exogenous 5-HTP supplementation accelerates sevoflurane emergence in a dose-dependent manner

5-HTP, the direct blood–brain barrier-permeable precursor of 5-HT, elevates central serotonergic tone. We adopted multi-modal delivery strategies to characterize its pro-arousal effects across naive, CRS and LPS mice (Fig. 1C). Repeated intraperitoneal (i.p.) 24 h interval 5-HTP injection produced no significant changes in anesthesia induction time at all tested doses (P > 0.05; Fig. 1D-F). In naive mice, 50 and 75 mg/kg 5-HTP significantly shortened emergence latency (P < 0.05, P < 0.01). Pathological mice displayed blunted sensitivity and required higher effective doses: CRS mice responded to 100 mg/kg, while LPS mice required 125 mg/kg 5-HTP to achieve comparable arousal facilitation (P < 0.05). To isolate central-specific effects, we performed intracerebroventricular (ICV) cannulation and drug delivery (Fig. 1G). ICV infusion of 22.7 mmol/L 5-HTP robustly reduced emergence time with no impact on induction (P < 0.001), whereas the low 11.35 mmol concentration exerted no behavioral effect. (Fig. 1H) We next localized the core arousal-relevant region to the central amygdala (CeA). Basal c-Fos neuronal activity in the CeA was markedly suppressed post-sevoflurane exposure compared with pre-anesthetic baseline (Fig. 1I-J). Local CeA microinjection of 2.5 or 5 mg/mL 5-HTP selectively shortened emergence time without altering induction duration (P < 0.01, P < 0.001; Fig. 1K-L). Collectively, systemic, central ventricular and region-specific CeA delivery of 5-HTP exert graded pro-arousal effects, with emotional disorder states desensitizing animals to peripheral serotonergic supplementation.

### 2.3 CeA serotonergic neuronal activity is suppressed under anxiety/depression and partially rescued by 5-HTP

We deployed TPH2-dependent fiber photometry to record real-time calcium dynamics of CeA 5-HT neurons across wakefulness, LORR, stable sevoflurane anesthesia and emergence phases (Fig. 2A-B). Bilateral CeA injection of rAAV-TPH2-GCaMP6S enabled cell-type-specific calcium signal readout via implanted optical fibers. In naive mice, CeA 5-HT neuron ΔF/F peak amplitudes were drastically attenuated during stable sevoflurane maintenance and rebounded during emergence (P < 0.05; Fig. 2C). This anesthetic suppression was further exacerbated in CRS and LPS mice, which exhibited persistently lower calcium peak signals throughout all recording stages (Fig. 2C-E). We combined i.p. 5-HTP administration with in vivo calcium imaging to test neuronal responsiveness to serotonergic augmentation (Fig. 2G-J). In naive animals, 75 mg/kg 5-HTP significantly elevated emergence-phase calcium transients of CeA 5-HT neurons (P < 0.05). In contrast, even elevated doses tailored for pathological models (100 mg/kg for CRS, 125 mg/kg for LPS) failed to fully restore blunted neuronal activity (P < 0.05; Fig. 2G-J). These fiber photometry results demonstrate that chronic anxiety and depression induce persistent hypoexcitability of CeA serotonergic neurons and impair their responsiveness to 5-HTP rescue.

**Figure 2.**
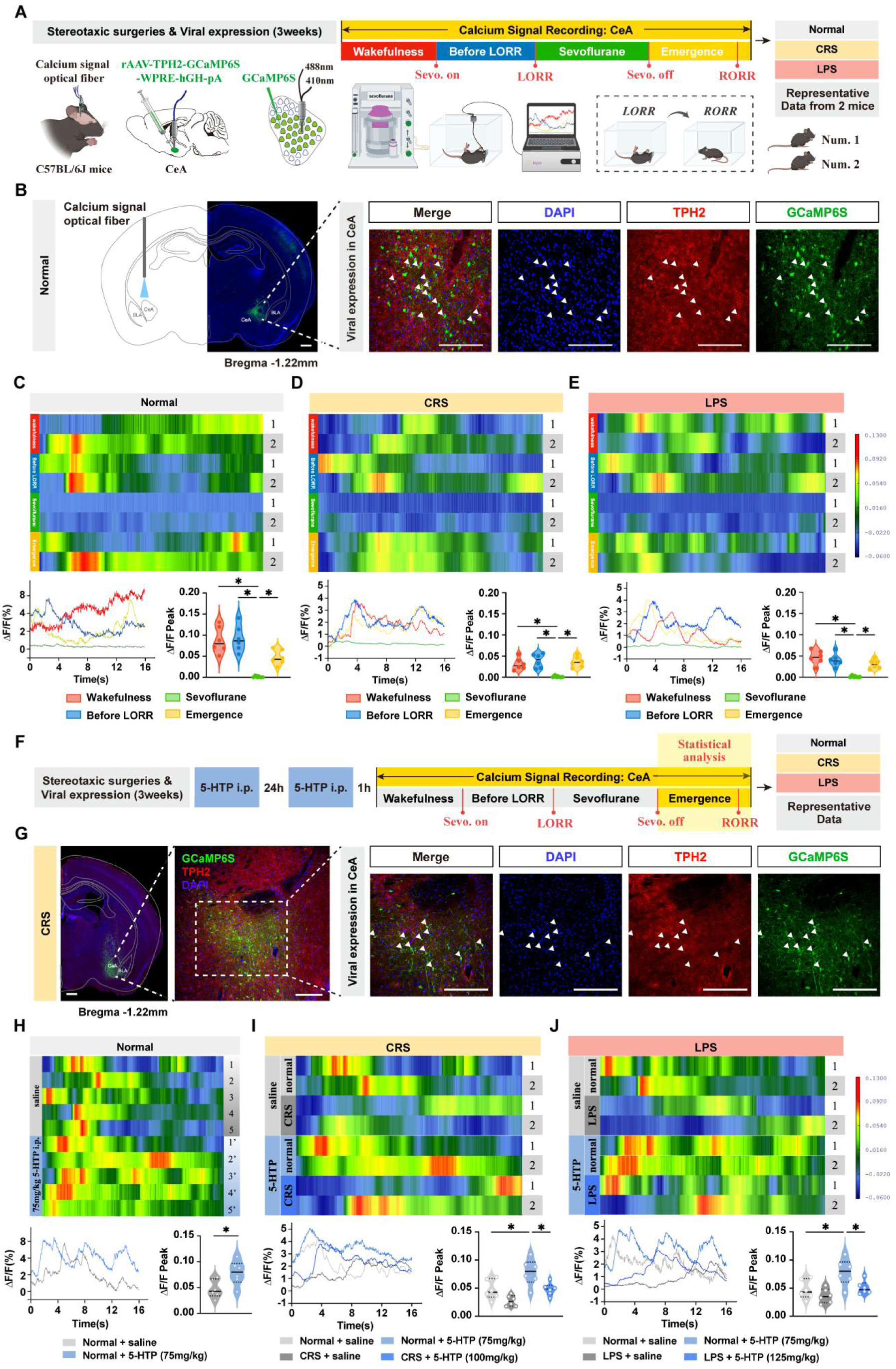
Fiber photometry recordings reveal that the neuronal activity of CeA^5-HT^ neurons is inhibited during sevoflurane anesthesia and shows blunted restoration by 5-HTP. **(A)** Experimental protocol for calcium signal recording of CeA^5-HT^ neurons. **(B)** Representative coronal brain slice of normal mice, showing the location of virus (rAAV-TPH2-GCaMP6S-WPRE-hGH-pA) injected in the CeA, and representative images of staining for DAPI, TPH2 and GCaMP6S in the CeA. **(C-E)** Heatmap and statistical diagram for show the activity wave of calcium signals and the peak of ΔF/F in the CeA of normal, CRS and LPS mice during the Wakefulness, Before LORR, Sevoflurane and Emergence four stages. **(F)** Experimental protocol for calcium signal recording of CeA^5-HT^ neurons and i.p. injection in the normal + saline, CRS + saline, LPS + saline, normal + 75 mg/kg 5-HTP, CRS + 100 mg/kg 5-HTP and LPS + 125 mg/kg 5-HTP groups during the emergence stage. **(G)** Representative coronal brain slice of CRS mice, showing the location of virus (rAAV-TPH2-GCaMP6S-WPRE-hGH-pA) injected in the CeA, and representative images of staining for DAPI, TPH2 and GCaMP6S in the CeA. **(H)** Heatmap and statistical diagram show the activity wave of calcium signals and the peak of ΔF/F for i.p. injection in the saline control and 75 mg/kg 5-HTP groups in the CeA of normal mice during the emergence stage. **(I)** Heatmap and statistical diagram show the activity wave of calcium signals and the peak of ΔF/F for i.p. injection in the normal + saline, CRS + saline, normal + 75 mg/kg 5-HTP and CRS + 100 mg/kg 5-HTP groups in the CeA of normal and CRS mice during the emergence stage. **(J)** Heatmap and statistical diagram show the activity wave of calcium signals and the peak of ΔF/F for i.p. injection in the normal + saline, LPS + saline, normal + 75 mg/kg 5-HTP and LPS + 125 mg/kg 5-HTP groups in the CeA of normal and LPS mice during the emergence stage. *p < 0.05; Scale bar: 100 μm; Data are mean ± SEM; i.p., intraperitoneal injection; 5-HTP, 5-hydroxytryptophan; CeA, central amygdala nucleus; Sevo, sevoflurane; LORR, loss of righting reflex; RORR, recovery of righting reflex; CRS, chronic restraint stress; LPS, lipopolysaccharide

### 2.4 Optogenetic and chemogenetic activation of CeA serotonergic neurons promotes arousal from sevoflurane anesthesia

We utilized TPH2 promoter-driven viral tools to achieve cell-type-specific optogenetic and chemogenetic manipulation of CeA 5-HT neurons (Fig. 3A). Stereotaxic delivery of pAAV-TPH2-ChETA-EYFP to the CeA enabled blue-light dependent neuronal excitation. Immunofluorescence quantification confirmed that 15 mW photostimulation robustly increased c-Fos/TPH2 co-labeling in the CeA relative to sham light conditions (P < 0.001; Fig. 3B-C). Optogenetic stimulation exerted no influence on induction time but shortened emergence latency in an intensity-dependent fashion in naive mice: 10 mW produced moderate effects (P < 0.05), while 15 mW yielded stronger arousal facilitation (P < 0.01). Unilateral and bilateral CeA photostimulation generated equivalent pro-arousal outcomes (P > 0.05; Fig. 3D). In both CRS and LPS mice, 15 mW CeA optogenetic activation significantly reversed delayed emergence (P < 0.01; Fig. 3E-F). For chemogenetic validation, rAAV-TPH2-hM3Dq was injected into the CeA to enable CNO-dependent neuronal excitation. Control experiments in virus-free mice confirmed that CNO alone did not alter anesthetic parameters (Fig. 3I). CNO administration elevated c-Fos expression in CeA 5-HT neurons of both naive and CRS animals, though activation magnitude was reduced in CRS mice (P < 0.01, P < 0.05; Fig. 3G-I). Chemogenetic activation dose-dependently accelerated emergence without changing induction time. Naive mice responded to low/intermediate CNO doses (0.25, 0.5 mg/kg), while CRS and LPS mice required higher 1.0 mg/kg CNO to achieve significant rescue (P < 0.01; Fig. 3J-L). We additionally performed chemogenetic inhibition via CeA rAAV-TPH2-hM4Di delivery (Supplementary Fig. 5C-G). CeA 5-HT neuronal suppression markedly prolonged emergence in naive mice (P < 0.01) but produced no behavioral shift in LPS animals, consistent with a pre-existing hypoactive baseline state of CeA serotonergic circuits under depressive pathology. To dissect upstream dorsal raphe (DR) contributions, we optogenetically activated DR-to-CeA serotonergic fibers (Supplementary Fig. 6A-B); 20 mW stimulation shortened emergence time (P < 0.05). Combined DR chemogenetic inhibition and CeA optogenetic activation assays further verified that CeA 5-HT neurons exert dominant arousal regulatory functions independent of upstream DR input (Supplementary Fig. 6C-D).

**Figure 3.**
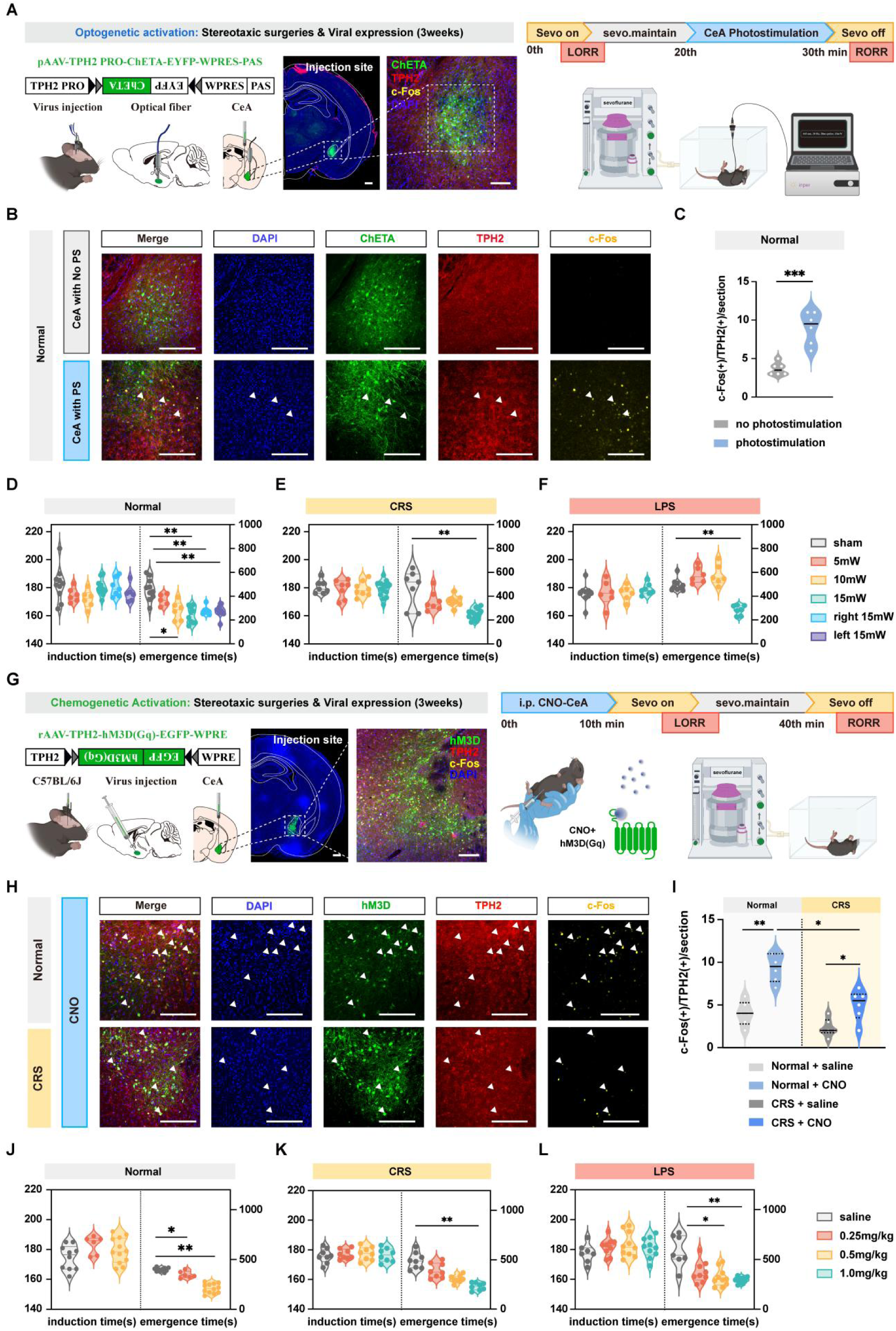
Optogenetic and chemogenetic activation of CeA^5-HT^ neurons promotes arousal from sevoflurane anesthesia in normal, CRS and LPS mice. **(A)** Experimental protocol for optogenetic activation of CeA^5-HT^ neurons. Schematic of the position of the virus injection and optic fiber implantation in the CeA, and representative coronal brain slice, showing the location of virus (pAAV-TPH2 PRO-ChETA-EYFP-WPRES-PAS) injection in the CeA. **(B)** Representative images of staining for c-Fos, TPH2, ChETA and DAPI in the CeA of normal mice with and without CeA photostimulation. **(C)** The quantification of c-Fos(+)/TPH2(+)/section in the CeA with or without CeA photostimulation. **(D)** Induction time and emergence time for normal mice in the sham, 5 mW, 10mW, 15 mW, right 15 mW and left 15 mW groups. **(E-F)** Induction time and emergence time for CRS and LPS mice in the sham, 5 mW, 10 mW and 15 mW groups. **(G)** Experimental protocol for chemogenetic activation of CeA^5-HT^ neurons. Schematic of the position of the virus injection in the CeA, and representative coronal brain slice, showing the location of virus (rAAV-TPH2-hM3D(Gq)- EGFP-WPRE) injection in the CeA. **(H)** Representative images show the co-expression of and staining for c-Fos, hM3D, TPH2 and DAPI in the CeA of normal and CRS mice with CNO activation. **(I)** The quantification of c-Fos(+)/TPH2(+)/section in the CeA for the Normal + saline, Normal + CNO, CRS + saline and CRS + CNO groups. **(J)** Induction time and emergence time for normal mice for i.p. injection in the saline control, 0.25 mg/kg CNO, 0.50 mg/kg CNO groups. **(K-L)** Induction time and emergence time for CRS and LPS mice for i.p. injection in the saline control, 0.25 mg/kg CNO, 0.50 mg/kg CNO and 1.0 mg/kg CNO groups. ***p < 0.001; **p < 0.01; *p < 0.05; Scale bar: 100 μm; Data are mean ± SEM; i.p., intraperitoneal injection; CNO, clozapine-N-oxide; CeA, central amygdala nucleus; Sevo, sevoflurane; LORR, loss of righting reflex; RORR, recovery of righting reflex; CRS, chronic restraint stress; LPS, lipopolysaccharide

To independently confirm the functional specificity of CeA serotonergic neurons, we generated and genotyped Sert-Cre mice (Supplementary Fig. 2). Sert-Cre founders on a C57BL/6J background were crossed with wild-type littermates for serial generational breeding, and PCR genotyping with sert-F/R1 (wild-type 295 bp) and sert-F/R2 (mutant 195 bp) primer pairs distinguished homozygous, heterozygous and wild-type offspring (Supplementary Fig. 2A-B). We injected Cre-dependent hChR2 virus into the CeA of Sert-Cre mice for optogenetic verification (Supplementary Fig. 4). Robust c-Fos induction was observed following 15 mW photostimulation of CeA Sert+ neurons (P < 0.001), and graded light stimulation recapitulated the dose-dependent pro-arousal phenotype observed in TPH2 viral experiments (Supplementary Fig. 4A-D). Parallel optogenetic activation of mPFC Sert-expressing neurons in Sert-Cre mice reproduced equivalent arousal-promoting effects (Supplementary Fig. 3A–D), cross-validating the conserved function of this serotonergic circuit across Cre driver lines.

### 2.5 Reciprocal serotonergic anatomical projections exist between CeA and mPFC

We combined retrograde cholera toxin subunit B (CTB-555) tracing and trans-monosynaptic rabies virus (RV) tracing to map CeA–medial prefrontal cortex (mPFC) serotonergic connectivity (Fig. 4A). CTB-555 microinjection into the mPFC retrogradely labeled abundant CeA neurons that co-expressed TPH2 (Fig. 4B-D). For monosynaptic tracing, a cocktail of TPH2-dependent helper viruses (rAAV-EF1α-DIO-TVA/oRVG + rAAV-TPH2-Cre) was delivered to the CeA, followed by EnvA-pseudotyped RV-EGFP injection three weeks later (Fig. 4E-G). One week post-RV infusion, numerous RV-labeled TPH2-positive serotonergic neurons were detected within the mPFC (Fig. 4H). Together, these anatomical tracing data establish a direct bidirectional 5-HTergic projection circuit linking the CeA and mPFC.

**Figure 4.**
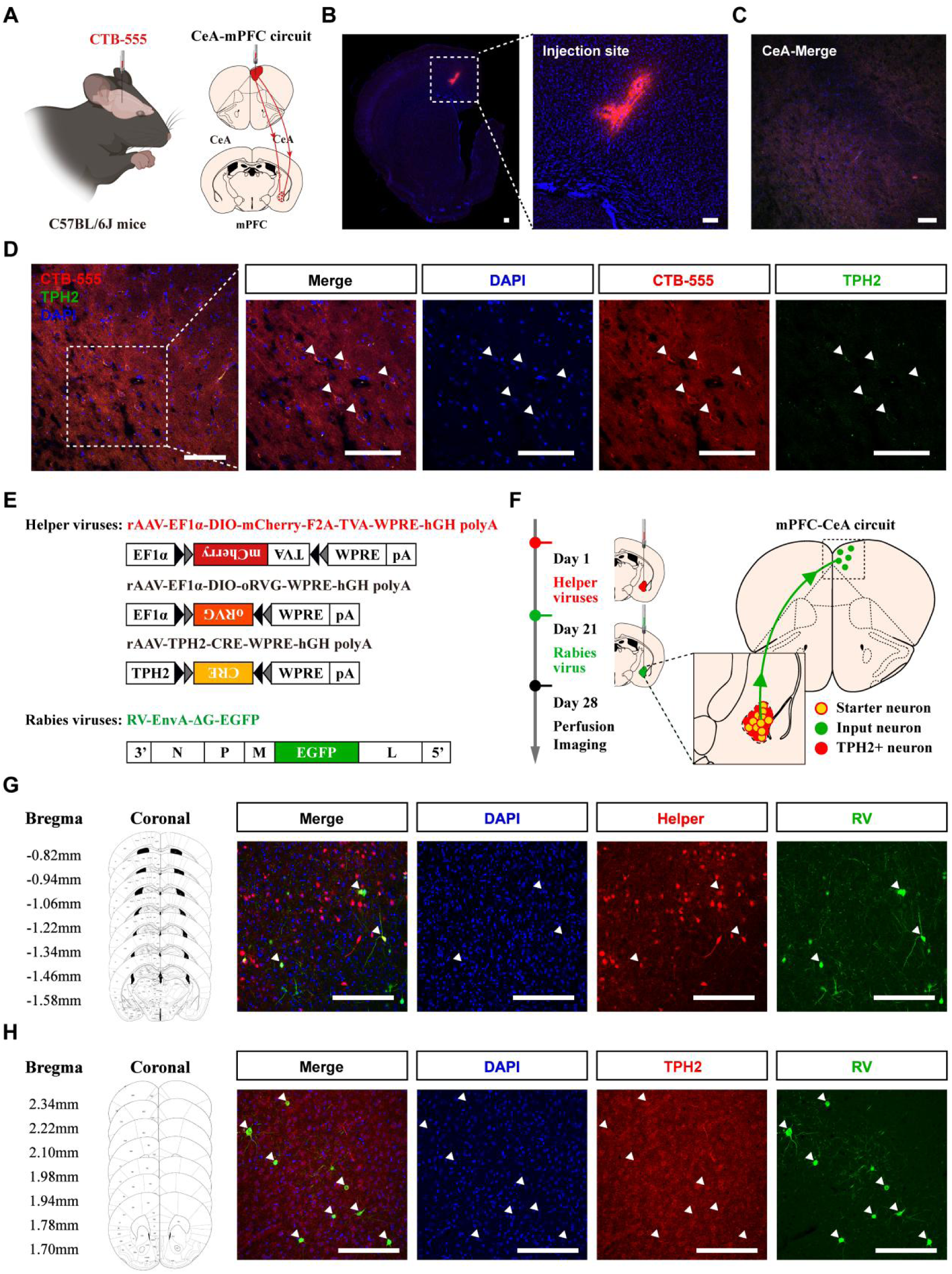
The existence of 5-HTergic bidirectional projection relationship between the CeA and mPFC. **(A)** Schematic of the injection position of retrograde labeling CTB-555. **(B)** Representative coronal brain slice of retrograde labeling. **(C-D)** Representative images show the co-expression of and staining for CTB and TPH2 in the CeA. **(E)** Design of viral vectors for RV-based trans-monosynaptic retrograde tracing, including helper viruses with cre-dependent expression of TVA receptor (rAAV-EF1a-DIO-mCherry-F2A-TVA-WPRE-hGH polyA), oRvG (rAAV-EF1⍺-DIO-oRVG-WPRE-hGH polyA) and TPH2(rAAV-TPH2-CRE-WPRE-hGH polyA). The RV was genetically modified by pseudotyping with EnvA (RV-EnvA-ΔG-EGFP). **(F)** Schematic of the injection procedure and experimental protocol for the helper viruses and RV injection in the TPH2+ neuron. Coronal schematic of retrograde tracing of 5-HTergic neurons (starter neurons, yellow; rAAV helper labeled TPH2+ neurons, red; RV labeled neurons, green). **(G)** Representative images show the co-expression of and staining for RV, helper and DAPI in the CeA. **(H)** Representative images show the co-expression of and staining for RV, TPH2 and DAPI in the mPFC. Scale bar: 100 μm; CTB, cholera toxin subunit B; CeA, central amygdala nucleus; mPFC, medial prefrontal cortex; TVA, tumor virus A; oRvG, optimized rabies virus glycoprotein; TPH2, tryptophan hydroxylase 2; RV, rabies virus

### 2.6 mPFC serotonergic neuronal activity is suppressed under anxiety/depression and weakly responsive to 5-HTP rescue

Given the reciprocal CeA–mPFC serotonergic anatomy, we next characterized mPFC 5-HT neuron dynamics and pharmacological responsiveness (Fig. 5A). Local mPFC microinjection of 2.5, 5 or 10 mg/mL 5-HTP significantly shortened emergence time with no alteration of induction latency (P < 0.05, P < 0.01, P < 0.001; Fig. 5B). Fiber photometry recording of mPFC TPH2-GCaMP6s signals followed identical experimental staging protocols used for CeA recordings (Fig. 5A). In naive mice, mPFC 5-HT calcium peaks were strongly suppressed during stable sevoflurane anesthesia; this inhibitory phenotype was exacerbated in CRS and LPS mice across all recording windows (P < 0.05; Fig. 5C–E). We further tested the impact of systemic 5-HTP on mPFC neuronal calcium activity during emergence (Fig. 5H-I). In naive animals, 75 mg/kg i.p. 5-HTP robustly amplified emergence-phase ΔF/F transients (P < 0.05). In contrast, maximal model-tailored 5-HTP doses failed to restore blunted mPFC serotonergic calcium signals in CRS (100 mg/kg) and LPS (125 mg/kg) mice (P < 0.05; Fig. 5J-L). These results mirror our CeA recordings and demonstrate widespread desensitization of the entire CeA–mPFC serotonergic network in pathological emotional states.

**Figure 5.**
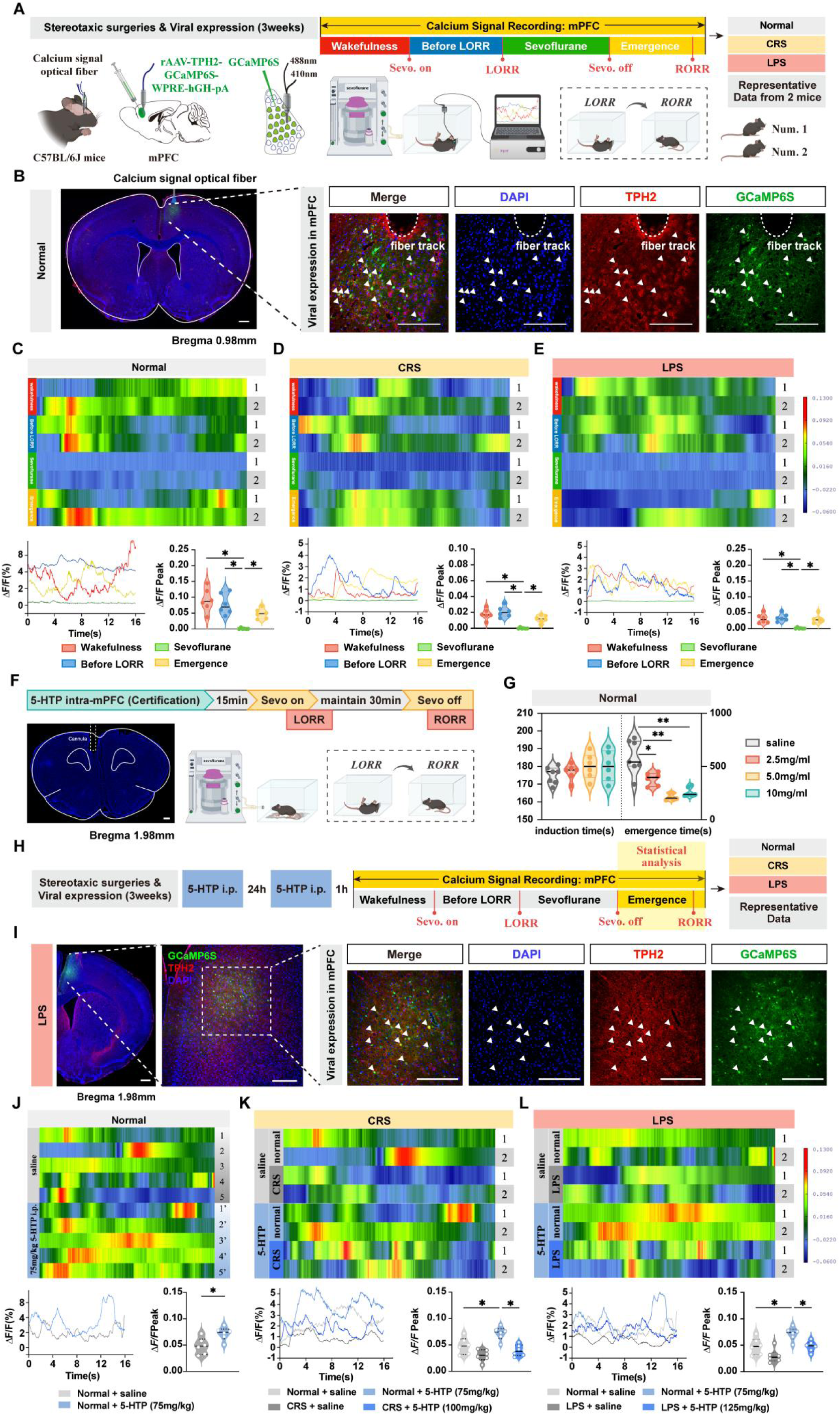
Fiber photometry recordings reveal that the neuronal activity of mPFC^5-HT^ neurons is inhibited during sevoflurane anesthesia and shows blunted restoration by 5-HTP. **(A)** Experimental protocol for calcium signal recording of mPFC^5-HT^ neurons. **(B)** Representative coronal brain slice of normal mice, showing the location of virus (rAAV-TPH2-GCaMP6S-WPRE-hGH-pA) injected in the mPFC, and representative images of staining for DAPI, TPH2 and GCaMP6S in the mPFC. **(C-E)** Heatmap and statistical diagram for show the activity wave of calcium signals and the peak of ΔF/F in the mPFC of normal, CRS and LPS mice during the Wakefulness, Before LORR, Sevoflurane and Emergence four stages. **(F)** Experimental protocol for intra-mPFC microinjection of different doses of 5-HTP, and representative image of the track of cannula implanted into the mPFC. **(G)** Induction time and emergence time for normal mice for intra-mPFC microinjection in the saline control, 2.5 mg/ml 5-HTP, 5.0 mg/ml 5-HTP and 10 mg/ml 5-HTP groups. **(H)** Experimental protocol for calcium signal recording of mPFC^5-HT^ neurons and i.p. injection in the normal + saline, CRS + saline, LPS + saline, normal + 75 mg/kg 5-HTP, CRS + 100 mg/kg 5-HTP and LPS + 125 mg/kg 5-HTP groups during the emergence stage. **(I)** Representative coronal brain slice of LPS mice, showing the location of virus (rAAV-TPH2-GCaMP6S-WPRE-hGH-pA) injected in the mPFC, and representative images of staining for DAPI, TPH2 and GCaMP6S in the mPFC. **(J)** Heatmap and statistical diagram show the activity wave of calcium signals and the peak of ΔF/F for i.p. injection in the saline control and 75 mg/kg 5-HTP groups in the mPFC of normal mice during the emergence stage. **(K)** Heatmap and statistical diagram show the activity wave of calcium signals and the peak of ΔF/F for i.p. injection in the normal + saline, CRS + saline, normal + 75 mg/kg 5-HTP and CRS + 100 mg/kg 5-HTP groups in the mPFC of normal and CRS mice during the emergence stage. **(L)** Heatmap and statistical diagram show the activity wave of calcium signals and the peak of ΔF/F for i.p. injection in the normal + saline, LPS + saline, normal + 75 mg/kg 5-HTP and LPS + 125 mg/kg 5-HTP groups in the mPFC of normal and LPS mice during the emergence stage. *p < 0.05; Scale bar: 100 μm; Data are mean ± SEM; i.p., intraperitoneal injection; 5-HTP, 5-hydroxytryptophan; mPFC, medial prefrontal cortex; Sevo, sevoflurane; LORR, loss of righting reflex; RORR, recovery of righting reflex; CRS, chronic restraint stress; LPS, lipopolysaccharide

### 2.7 Optogenetic and chemogenetic activation of mPFC serotonergic neurons facilitates sevoflurane arousal

We performed cell-type-specific optogenetic modulation of mPFC 5-HT neurons using TPH2-ChETA viral delivery (Fig. 6A). 15 mW photostimulation efficiently elevated c-Fos/TPH2 co-expression in the mPFC (Fig. 5B). Optogenetic stimulation produced no shift in induction time, yet dose-dependently accelerated emergence in naive mice: 10 mW exerted mild effects (P < 0.05), while 15 mW generated robust facilitation (P < 0.001). CRS and LPS mice required elevated 20 mW light intensity to achieve statistically significant arousal rescue (P < 0.05; Fig. 5C-E). Chemogenetic activation via mPFC TPH2-hM3Dq delivery yielded consistent behavioral outcomes (Fig. 5F). CNO treatment increased c-Fos expression in mPFC 5-HT neurons of naive and LPS mice, though activation magnitude remained lower in LPS animals (P < 0.001, P < 0.05; Fig. 5G-H). Naive mice responded to 1.0 mg/kg CNO, whereas CRS/LPS cohorts required 2.0 mg/kg CNO for comparable emergence shortening (P < 0.05; Fig. 5I-K). Parallel optogenetic assays in Sert-Cre mice selectively targeting mPFC serotonergic neurons recapitulated these dose-dependent pro-arousal phenotypes (Supplementary Fig. 3), independently validating mPFC 5-HT neuron function in consciousness recovery. Combined optogenetic and chemogenetic evidence from both TPH2 viral and Sert-Cre transgenic models confirms that mPFC serotonergic activation accelerates anesthetic arousal, with pathological states reducing neuronal responsiveness to stimulation.

**Figure 6.**
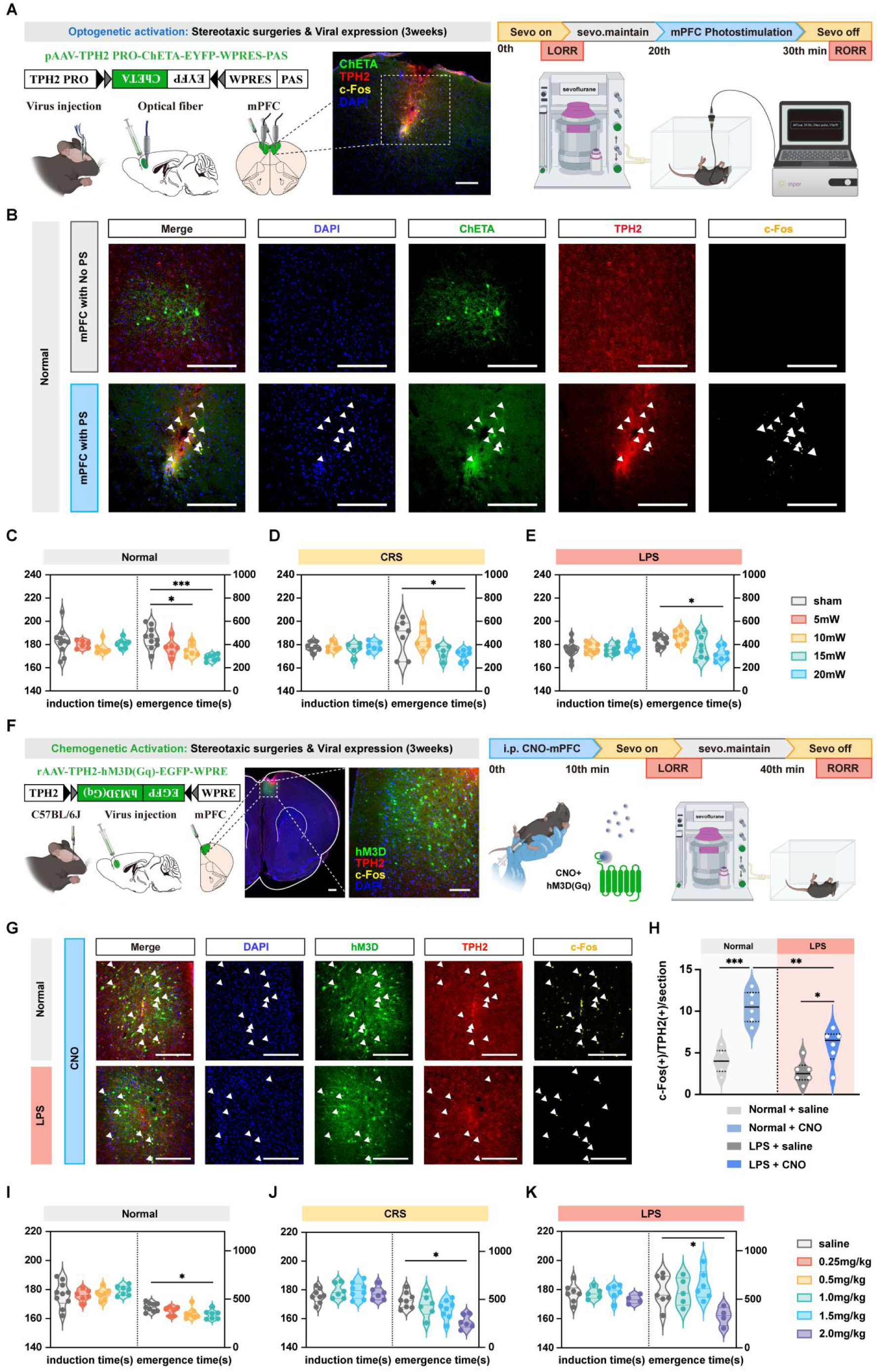
Optogenetic and chemogenetic activation of mPFC^5-HT^ neurons reduced the emergence time of sevoflurane anesthesia in normal, CRS and LPS mice. **(A)** Experimental protocol for optogenetic activation of mPFC^5-HT^ neurons. Schematic of the position of the virus injection and optic fibers implantation in the mPFC, and representative coronal brain slice, showing the location of virus (pAAV-TPH2 PRO-ChETA-EYFP-WPRES-PAS) injection in the mPFC. **(B)** Representative images of staining for c-Fos, TPH2, ChETA and DAPI in the mPFC of normal mice with and without mPFC photostimulation. **(C)** Induction time and emergence time for normal mice in the sham, 5 mW, 10 mW and 15 mW groups. **(D)** Induction time and emergence time for CRS mice in the sham, 10 mW, 15 mW and 20 mW groups. **(E)** Induction time and emergence time for LPS mice in the sham, 10 mW, 15 mW and 20 mW groups. **(F)** Experimental protocol for chemogenetic activation of mPFC^5-HT^ neurons. Schematic of the position of the virus injection in the mPFC, and representative coronal brain slice, showing the location of virus (rAAV-TPH2-hM3D(Gq)- EGFP-WPRE) injection in the mPFC. **(G)** Representative images show the co-expression of and staining for c-Fos, hM3D, TPH2 and DAPI in the mPFC of normal and LPS mice with CNO activation. **(H)** The quantification of c-Fos(+)/TPH2(+)/section in the mPFC for the Normal + saline, Normal + CNO, LPS + saline and LPS + CNO groups. **(I)** Induction time and emergence time for normal mice for i.p. injection in the saline control, 0.25 mg/kg CNO, 0.50 mg/kg CNO and 1.0 mg/kg CNO groups. **(J)** Induction time and emergence time for CRS mice for i.p. injection in the saline control, 1.0 mg/kg CNO, 1.50 mg/kg CNO and 2.0 mg/kg CNO groups. **(K)** Induction time and emergence time for LPS mice for i.p. injection in the saline control, 1.0 mg/kg CNO, 1.50 mg/kg CNO and 2.0 mg/kg CNO groups. ***p < 0.001; **p < 0.01; *p < 0.05; Scale bar: 100 μm; Data are mean ± SEM; i.p., intraperitoneal injection; CNO, clozapine-N-oxide; mPFC, medial prefrontal cortex; Sevo, sevoflurane; LORR, loss of righting reflex; RORR, recovery of righting reflex; CRS, chronic restraint stress; LPS, lipopolysaccharide

### 2.8 Bidirectional 5-HT1A and 5-HT2A receptor signaling mediates CeA–mPFC circuit arousal function

We combined regional receptor antagonism with circuit-specific neuronal activation to identify the downstream molecular receptors transducing pro-arousal serotonergic signals (Fig. 10). First, we tested mPFC-localized 5-HT1A (WAY-100635) and 5-HT2A (KET) blockade against CeA 5-HT neuron activation. CeA microinjection of 5 mg/mL 5-HTP shortened emergence, an effect fully abolished by intra-mPFC WAY infusion (P < 0.01; Fig. 7A-B). When paired with CeA optogenetic activation, mPFC delivery of either WAY or KET completely ablated the arousal-facilitating effect across naive, CRS and LPS mice (Fig. 7C-F). Chemogenetic CeA activation produced identical rescue phenotypes that were reversed via mPFC microinjection of the two receptor antagonists; notably, KET lost inhibitory efficacy in pathological CRS/LPS groups, while WAY retained full blocking capacity (Fig. 7D–J). Reciprocal pharmacological experiments targeted CeA receptors during mPFC serotonergic activation (Fig. 7L-R). Microinjection of WAY or KET into the CeA fully eliminated the pro-arousal behavioral shifts induced by mPFC optogenetic and chemogenetic stimulation in all three experimental cohorts (Fig. 7L-R). Collectively, bidirectional pharmacological perturbation demonstrates that both mPFC and CeA-expressed 5-HT1A and 5-HT2A receptors are indispensable downstream mediators of CeA–mPFC serotonergic circuit-dependent anesthetic arousal.

**Figure 7.**
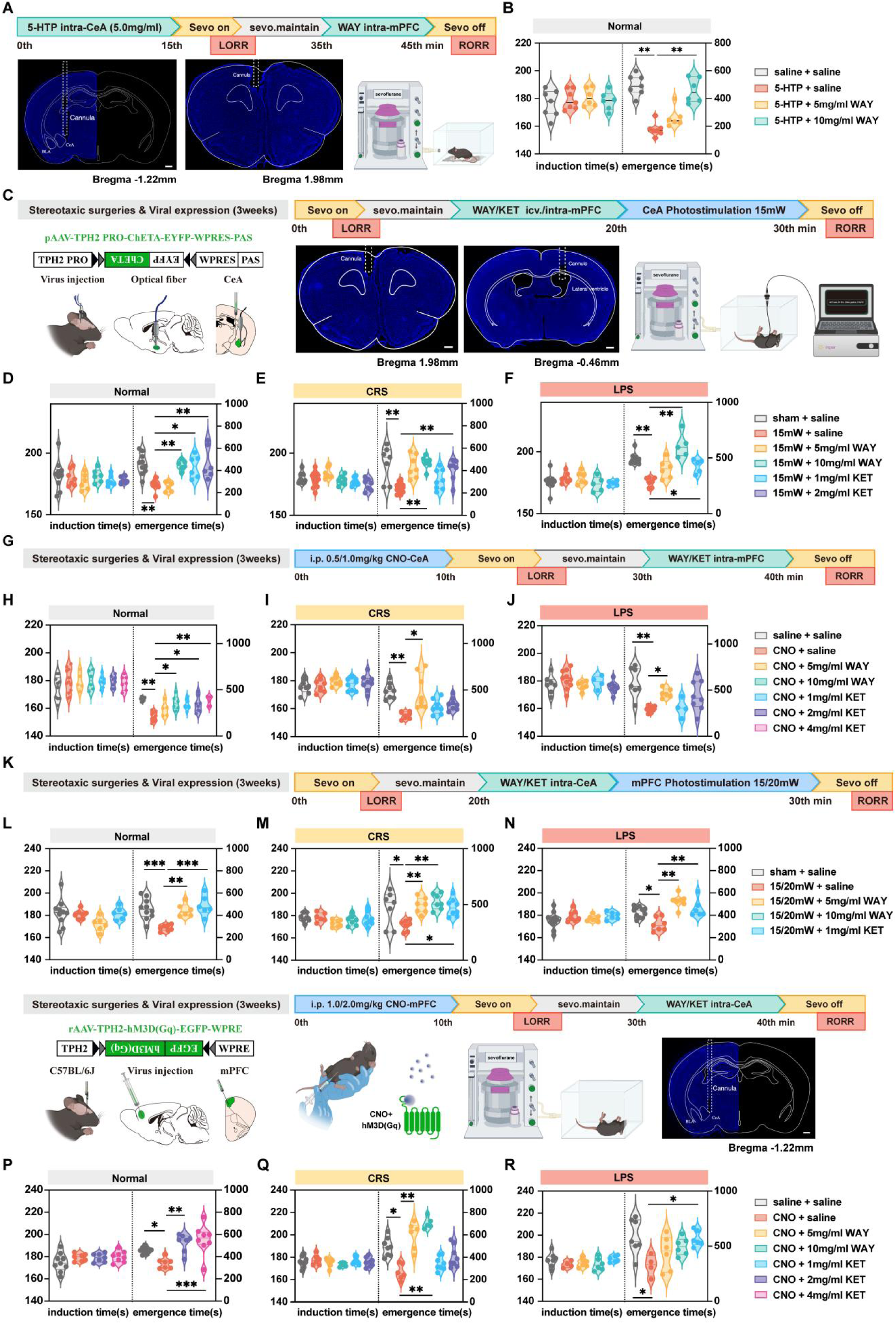
Optogenetic and chemogenetic activation combined with pharmacological blockade of 5-HT1AR and 5-HT2AR reveals bidirectional 5-HTergic interactions between the CeA and mPFC. **(A)** Experimental protocol for intra-CeA microinjection of 5.0 mg/ml 5-HTP and intra-mPFC microinjection of 5-HT1AR antagonist WAY and representative image of the track of cannula implanted into the CeA and mPFC. **(B)** Induction time and emergence time for normal mice in the saline + saline, 5-HTP + saline, 5-HTP + 5.0 mg/ml WAY and 5-HTP + 10 mg/ml WAY groups. **(C)** Experimental protocol for optogenetic activation of CeA^5-HT^ neurons (15 mW) and icv. injection of 5-HT1AR antagonist WAY, and intra-mPFC microinjection of 5-HT2AR antagonist KET. Representative image of the track of cannula implanted into the mPFC and lateral ventricle. **(D, E)** Induction time and emergence time for normal mice and CRS mice in the sham + saline, 5 mW + saline, 15 mW + 5 mg/ml WAY, 15 mW + 10 mg/ml WAY, 15 mW + 1 mg/ml KET and 15 mW + 2 mg/ml KET groups. **(F)** Induction time and emergence time for LPS mice in the sham + saline, 5 mW + saline, 15 mW + 5 mg/ml WAY, 15 mW + 10 mg/ml WAY and 15 mW + 1 mg/ml KET groups. **(G)** Experimental protocol for chemogenetic activation of CeA^5-HT^ neurons and microinjection of WAY or KET into the mPFC, and CNO was administered at 0.5 mg/kg in the normal mice and 1.0 mg/kg in both the CRS and LPS mice. **(H)** Induction time and emergence time for normal mice in the saline + saline, CNO + saline, CNO + 5 mg/ml WAY, CNO + 10 mg/ml WAY, CNO + 1 mg/ml KET, CNO + 2 mg/ml KET and CNO + 4 mg/ml KET groups. **(I, J)** Induction time and emergence time for CRS and LPS mice in the saline + saline, CNO + saline, CNO + 5 mg/ml WAY, CNO + 1 mg/ml KET and CNO + 2 mg/ml KET. **(K)** Experimental protocol for optogenetic activation of mPFC^5-HT^ neurons at 15 mW for the normal mice and 20 mW for the CRS and LPS mice, combined with microinjection of WAY or KET into the CeA. **(L)** Induction time and emergence time for normal mice in the sham + saline, 15 mW + saline, 15 mW + 5 mg/ml WAY and 15 mW + 1 mg/ml KET groups. **(M)** Induction time and emergence time for CRS mice in the sham + saline, 20 mW + saline, 20 mW + 5 mg/ml WAY, 20 mW + 10 mg/ml WAY and 20 mW + 1 mg/ml KET groups. **(N)** Induction time and emergence time for LPS mice in the sham + saline, 20 mW + saline, 20 mW + 5 mg/ml WAY and 20 mW + 1 mg/ml KET groups. **(O)** Experimental protocol for chemogenetic activation of mPFC^5-HT^ neurons and microinjection of WAY or KET into the CeA, and CNO was administered at 1.0 mg/kg in the normal mice and 2.0 mg/kg in both the CRS and LPS mice. Representative image of the track of cannula implanted into the CeA. **(P)** Induction time and emergence time for normal mice in the saline + saline, CNO + saline, CNO + 2 mg/ml KET and CNO + 4 mg/ml KET groups. **(Q)** Induction time and emergence time for CRS mice in the saline + saline, CNO + saline, CNO + 5 mg/ml WAY, CNO + 10 mg/ml WAY, CNO + 1 mg/ml KET and CNO + 2 mg/ml KET groups. **(R)** Induction time and emergence time for LPS mice in the saline + saline, CNO + saline, CNO + 5 mg/ml WAY, CNO + 10 mg/ml WAY and CNO + 1 mg/ml KET groups. ***p < 0.001; **p < 0.01; *p < 0.05; Scale bar: 100 μm; Data are mean ± SEM; i.p., intraperitoneal injection; icv., intracerebroventricular injection; 5-HTP, 5-hydroxytryptophan; WAY, WAY-100635; KET, ketanserin; CeA, central amygdala nucleus; mPFC, medial prefrontal cortex; Sevo, sevoflurane; LORR, loss of righting reflex; RORR, recovery of righting reflex; CRS, chronic restraint stress; LPS, lipopolysaccharide; 5-HT1AR, 5-HT1A receptor; 5-HT2AR, 5-HT2A receptor

### 2.9 Graded intervention intensity is required to rescue blunted serotonergic circuit function under emotional pathology

Correlation regression analyses quantified the relationship between intervention strength and emergence latency across naive, CRS and LPS mice (Fig. 8). In wild-type animals, emergence time exhibited strong negative linear correlations with increasing 5-HTP dosage, optogenetic light power and CNO chemogenetic dose: stronger stimulation yielded faster recovery of consciousness (Fig. 8A, E, I). These inverse linear correlations were entirely lost in CRS and LPS mice (Fig. 8B–C, J–K). Low-to-moderate interventions failed to produce arousal benefits and even exerted mild inhibitory effects on emergence in pathological animals: 75 mg/kg 5-HTP, 10 mW light and intermediate CNO doses all prolonged recovery time in CRS/LPS groups (Fig. 8D, H, L). These data indicate that anxiety and depression remodel serotonergic receptor sensitivity and intracellular downstream signaling cascades within the CeA–mPFC circuit, driving neuronal desensitization that necessitates escalated, high-intensity targeted interventions to reverse delayed anesthetic emergence.

**Figure 8.**
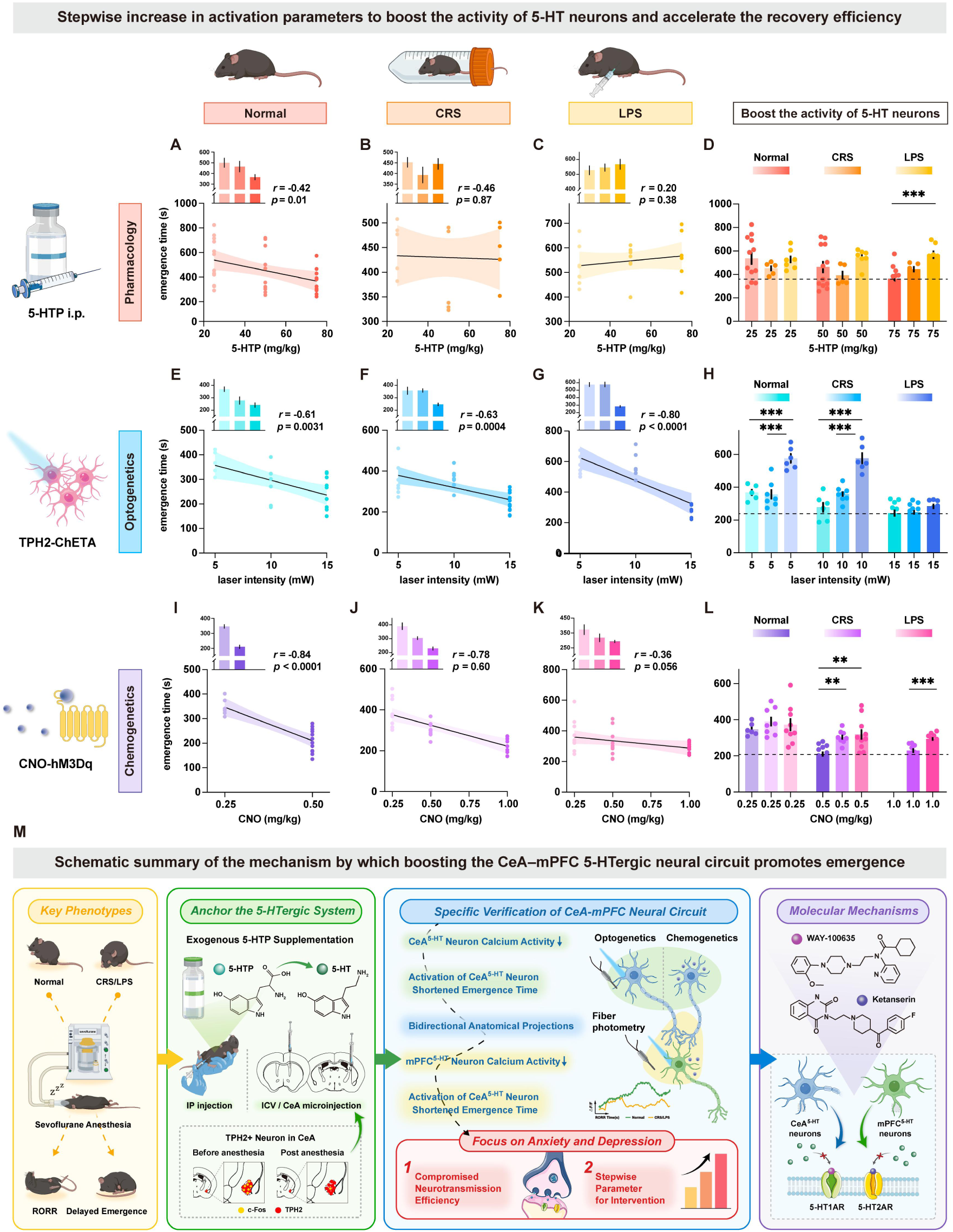
Stepwise increase in activation parameters using pharmacological, optogenetic, and chemogenetic approaches to boost the activity of 5-HT neurons and accelerate the emergence efficiency after sevoflurane anesthesia. **(A-D)** Effects of pharmacological activation of the 5-HTergic system on emergence from sevoflurane anesthesia. Correlation analyses between different doses of 5-HTP and emergence time are shown for normal mice **(A)**, CRS mice **(B)**, and LPS mice **(C)**, with corresponding emergence time after i.p. injection of 25, 50, or 75 mg/kg 5-HTP shown in the upper panels and correlation analyses shown in the lower panels. Emergence time after 5-HTP administration were further compared among the normal, CRS, and LPS groups **(D)**. **(E-H)** Effects of optogenetic activation of CeA^5-HT^ neurons on emergence from sevoflurane anesthesia. Correlation analyses between different laser intensities and emergence time are shown for normal mice **(E)**, CRS mice **(F)**, and LPS mice **(G)**, with corresponding emergence time under 5, 10, or 15 mW stimulation shown in the upper panels and correlation analyses shown in the lower panels. Emergence time after optogenetic activation of CeA^5-HT^ neurons were further compared among the normal, CRS, and LPS groups (H). **(I-L)** Effects of chemogenetic activation of CeA^5-HT^ neurons on emergence from sevoflurane anesthesia. Correlation analyses between different doses of CNO and emergence time are shown for normal mice **(I)**, CRS mice **(J)**, and LPS mice **(K)**, with corresponding emergence time after i.p. injection of 0.25, 0.5, or 1.0 mg/kg CNO shown in the upper panels and correlation analyses shown in the lower panels. Emergence time after CNO administration were further compared among the normal, CRS, and LPS groups **(L)**. **(M)** Schematic summary illustrating the proposed mechanism by which boosting the CeA-mPFC 5-HTergic neural circuit promotes emergence from sevoflurane anesthesia and facilitates recovery of consciousness under anxiety- and depression-like states. Anxiety- and depression-like conditions delayed emergence from sevoflurane anesthesia and were associated with reduced responsiveness of the 5-HTergic system. Pharmacological supplementation with 5-HTP, together with optogenetic and chemogenetic activation of CeA^5-HT^ neurons projecting to the mPFC, accelerated emergence. Bidirectional circuit-level validation further identified 5-HT1AR and 5-HT2AR as key mediators of the pro-emergence effect, as pharmacological blockade of these receptors abolished the behavioral effects. These findings suggest that anxiety- and depression-like states require stepwise parameter-optimized enhancement of 5-HTergic activity to effectively restore arousal from sevoflurane anesthesia. ***p < 0.001, **p < 0.01, Data are mean ± SEM; i.p., intraperitoneal injection; CRS, chronic restraint stress; LPS, lipopolysaccharide; 5-HTP, 5-hydroxytryptophan; CNO, clozapine-N-oxide; CeA, central amygdala nucleus; mPFC, medial prefrontal cortex; 5-HT1AR, 5-HT1A receptor; 5-HT2AR, 5-HT2A receptor

### 2.10 Summary of circuit functional remodeling under anxiety and depression

Multi-modal pharmacological, fiber photometry, optogenetic, chemogenetic and viral tracing experiments, plus independent validation in Sert-Cre transgenic mice, collectively define a core bidirectional CeA-mPFC serotonergic circuit governing sevoflurane arousal. Under physiological conditions, activation of this circuit enhances neuronal excitability and accelerates consciousness restoration. Chronic anxiety and depression drive persistent hypoactivity of CeA/mPFC 5-HT neurons and blunt 5-HT1A/5-HT2A receptor-mediated synaptic transmission, which mechanistically explains the clinical phenotype of prolonged anesthetic emergence in emotionally vulnerable subjects. Escalated circuit-specific stimulation targeting these two receptor subtypes effectively rescues pathological arousal deficits, establishing this reciprocal serotonergic pathway as a critical therapeutic target for perioperative anesthetic management in patients with anxiety and depressive comorbidities(Fig. 8M).

## 3 Discussion

Since consciousness is modulated by multiple neural factors, prolonged emergence from general anesthesia poses substantial clinical risks. Therefore, elucidating the regulatory mechanisms governing anesthetic recovery remains an urgent and important research topic. Comorbid anxiety and depression have become a global public health crisis^1^, and accumulating clinical evidence demonstrates that preoperative emotional disorders are tightly associated with delayed anesthetic emergence and severe postoperative complications. These adverse outcomes are largely attributed to disrupted neural circuits that regulate the sleep–wake cycle^3,4^. As one of the most widely used inhaled anesthetics, sevoflurane occupies a core position in clinical surgery and diagnostic procedures. A thorough dissection of the neural mechanisms underlying consciousness transitions during sevoflurane anesthesia not only provides direct guidance for optimizing perioperative anesthetic management, but also establishes a fundamental theoretical framework for understanding the general neurobiological rules of consciousness regulation.

As an essential monoamine neurotransmitter in the central nervous system, 5-HT participates extensively in consciousness-related physiological functions, and numerous studies have confirmed its indispensable role in sleep–wake modulation^42^. Our previous work revealed that activating 5-HTergic neurons in the DR and their projections to the BLA effectively shortens sevoflurane emergence time. Both presynaptic 5-HT₁A autoreceptors in the DR and postsynaptic 5-HT₁A receptors in the BLA mediate this pro-arousal effect^20,41^. Building on these findings, the present study further explores the neural circuit mechanisms by which the 5-HTergic system regulates consciousness recovery during sevoflurane anesthesia.

We first established mouse models with normal status or anxiety/depression-like phenotypes induced by CRS and LPS, and took the impact of preoperative emotional disorders on anesthetic emergence as the entry point. Behavioral results showed that mice with anxiety-and depression-like behaviors exhibited significantly prolonged emergence. This pathological phenotype indicates that clarifying the aberrant neural circuit mechanisms underlying delayed emergence under emotional disturbance can, in turn, help reveal the physiological regulatory rules of anesthetic recovery. By systematically comparing dynamic circuit activities across normal, CRS and LPS groups, we aimed to identify the core regulatory nodes for consciousness transitions.

CeA is a key component of the ascending reticular activating system and plays a vital role in initiating and maintaining wakefulness^43,44^. Previous studies have confirmed extensive reciprocal anatomical projections between the CeA and mPFC, both of which are abundantly distributed with 5-HT receptors. In addition, neuronal inactivation in the mPFC delays anesthetic emergence, and cerebral 5-HT levels decline markedly under anesthetic states^38,35^. Accordingly, we focused on characterizing how the CeA–mPFC 5-HTergic circuit and related receptors modulate the process of anesthetic emergence.

Pharmacological experiments showed that systemic, intracerebroventricular or local CeA supplementation of exogenous 5-HT notably shortened emergence duration. Fiber photometry recordings revealed that CeA 5-HTergic neuronal activity was suppressed during sevoflurane anesthesia and gradually recovered during the emergence phase. Selective activation of CeA 5-HTergic neurons via optogenetics and chemogenetics accelerated anesthetic emergence, which was consistent with the pro-arousal effect induced by 5-HTP. Given the bidirectional anatomical connections between the CeA and mPFC and the dynamic calcium signals recorded in the mPFC, we further verified that targeted activation of mPFC 5-HTergic neurons also facilitated anesthetic recovery. Of note, the optogenetic light intensity and chemogenetic CNO dosage required to activate mPFC neurons were much higher than those for CeA neurons, suggesting regional heterogeneity in the density and intrinsic activity of 5-HTergic neurons across distinct brain nuclei.

To rigorously confirm the specificity and functional reliability of the CeA–mPFC 5-HTergic circuit in regulating anesthetic emergence, we successfully bred TPH2-Cre transgenic mice and performed a series of complementary anatomical, cellular and functional validations from multiple dimensions, which further consolidated our core conclusions. The relevant evidence is elaborated as follows:

First, we conducted immunofluorescence colocalization assays to verify the cellular identity of recorded neurons in the mPFC. The results showed that the calcium indicator carried by fiber photometry viruses exhibited high colocalization with TPH2-positive serotonergic neurons in the mPFC at the cellular level. This finding confirms that the calcium signals we recorded primarily originated from TPH2-expressing 5-HTergic neurons, rather than other cell types.

Second, we combined **c-Fos** immunostaining to map neuronal activation during anesthetic emergence. Immunohistochemical results demonstrated that c-Fos was highly colocalized with TPH2-positive neurons in both the CeA and mPFC during the recovery phase from sevoflurane anesthesia. From the perspective of cellular activity, this evidence directly proves that TPH2-expressing serotonergic neurons are strongly activated during anesthetic emergence and serve as key cellular components driving the recovery of consciousness.

Third, we cross-validated the circuit function using **wild-type (WT)** mice and newly generated TPH2-Cre mice. Functional behavioral assays consistently confirmed that the CeA–mPFC 5-HTergic circuit is actively involved in promoting anesthetic emergence in both genotypes. The consistent phenotypic results across different mouse strains rule out the interference of genetic background and preliminarily verify the universal function of this circuit. Fourth, we adopted Cre-dependent DIO viral tools in TPH2-Cre mice for cell-type-specific manipulation. After stereotaxic injection of DIO viruses into the CeA and mPFC of TPH2-Cre mice, we observed high efficiency and specific infection of local TPH2-positive neurons. Subsequent behavioral tests further validated that specific modulation of these infected 5-HTergic neurons reliably altered sevoflurane emergence time. Combined with the above anatomical and cellular evidence, the DIO virus-based functional tests fully confirm the cell-type specificity of the CeA–mPFC 5-HTergic circuit in regulating anesthetic emergence. Collectively, multi-level evidence from immunofluorescence, neuronal activity labeling, cross-strain functional verification and Cre-dependent viral manipulation comprehensively demonstrates that the CeA–mPFC circuit exerts pro-arousal effects in a strict 5-HT neuron-dependent manner.

In terms of molecular mechanisms underlying circuit function, microinjection of the 5-HT₁A receptor antagonist WAY-100635 or 5-HT₂A receptor antagonist KET into the mPFC completely abolished the pro-arousal effect induced by optogenetic or chemogenetic activation of CeA 5-HTergic neurons. Similarly, intra-CeA injection of the two antagonists reversed the accelerated emergence caused by activating mPFC 5-HTergic neurons. Under anxiety- and depression-like conditions, specific activation of 5-HTergic neurons in the CeA or mPFC still effectively rescued delayed emergence, whereas local application of receptor antagonists eliminated this rescue effect. These results strongly indicate that bidirectional signaling transmission within the CeA-mPFC 5-HTergic circuit is central to anesthetic regulation, and synaptic transmission mediated by 5-HT₁A and 5-HT₂A receptors constitutes the core molecular mechanism.

Integrating behavioral performance and fiber photometry data, our study reveals that the CeA-mPFC 5-HTergic circuit modulates sevoflurane emergence mainly by regulating synaptic transmission efficiency. Under physiological conditions, exogenous 5-HT supplementation or artificial activation of this circuit effectively shortens emergence time. In contrast, a higher intensity of intervention is required to achieve equivalent pro-arousal effects in mice with anxiety- and depression-like phenotypes. Synaptic transmission efficiency depends on presynaptic neurotransmitter release, which is closely coupled with calcium influx and calcium signal activity in presynaptic neurons^46^. Calcium signal recordings showed that high-dose 5-HTP not only shortened emergence time but also enhanced neuronal calcium activity in model mice. Nevertheless, even high-dose 5-HTP failed to fully restore calcium signaling to normal levels in pathological states. This phenomenon suggests that anxiety and depression impair 5-HT receptor sensitivity and synaptic transmission efficiency^47,48^, which is the core pathological mechanism leading to delayed anesthetic emergence. Of note, presynaptic calcium dynamics and neurotransmitter release represent only the initial link of the regulatory cascade. They can trigger complex postsynaptic signaling events, including protein kinase cascades and transcriptional regulation, all of which may jointly facilitate the recovery of consciousness. The detailed mechanisms of these downstream pathways remain to be further explored.

The DR is the primary brain region generating 5-HTergic neurons and sends widespread projections to multiple brain areas including the CeA^49^. On the basis of our previous finding that the DR–BLA 5-HTergic circuit promotes anesthetic arousal^20,41^, we hypothesize that the CeA and mPFC act as secondary and tertiary functional nodes in a hierarchical 5-HTergic network to collectively regulate sevoflurane emergence. Further research is needed to clarify the complex regulatory relationships within this upstream-to-downstream circuit.

The present study is the first to reveal that synaptic transmission efficiency of the CeA– mPFC 5-HTergic circuit is critical for consciousness recovery after general anesthesia. Under anxiety and depression, the activity of this circuit is persistently suppressed, which directly explains the high susceptibility to delayed emergence in this population. Our findings confirm that potent activation of this specific serotonergic pathway can restore circuit function and reverse anesthetic recovery deficits. This work lays a solid mechanistic foundation for understanding anesthetic emergence in patients with emotional comorbidities (Fig. 12). In clinical practice, our results also provide feasible intervention strategies for preoperative patients with anxiety and depression, and offer theoretical support for future clinical cohort studies.

We also explain the data fluctuations observed in the original experiments: the emergence time of saline control groups across normal, CRS and LPS groups showed minor variations, which was mainly caused by ambient temperature differences (±0.5 °C) among experimental batches. Environmental temperature affects mouse metabolic rate and anesthetic clearance, and even subtle fluctuations can alter the recovery of the righting reflex, a common confounding factor in anesthetic emergence experiments. We have performed standardization and batch correction on all datasets during statistical analysis to eliminate such interference and ensure the reliability of experimental conclusions.

Apart from the circuit-targeted interventions proposed in this study, several alternative strategies with greater clinical translational potential are worthy of discussion. First, receptor-selective ligands: Compounds such as the 5-HT₁A agonist 8-OH-DPAT and 5-HT₂A agonist TCB-2 can specifically bind to corresponding receptors in the CeA–mPFC circuit. This approach achieves region-specific regulation and avoids the non-specific effects induced by global activation of the serotonergic system, thus improving intervention safety and efficacy. Second, neuromodulation techniques: Transcranial magnetic stimulation and deep brain stimulation enable precise modulation of the CeA–mPFC circuit via non-invasive or minimally invasive methods. These technologies are particularly suitable for patients with preoperative anxiety and depression who cannot tolerate pharmacological interventions, and have shown promising application prospects in regulating mental disorders and anesthetic recovery.

It is necessary to clearly define the research scope and avoid overinterpretation of conclusions. This study only focuses on the role of the CeA-mPFC 5-HTergic circuit in anesthetic emergence, rather than natural sleep-wake cycles. Although the regulatory networks for anesthetic recovery and natural sleep-wakefulness share partial similarities, they are essentially distinct physiological processes: anesthetic emergence refers to rapid consciousness recovery from drug-induced unconsciousness, while natural sleep–wake transition is a spontaneous physiological rhythm. Therefore, we revise our previous overstated description and clarify that the CeAmPFC serotonergic pathway is a key regulator of sevoflurane anesthetic emergence, rather than a core factor governing the natural sleep-wake cycle. In future work, we will combine **electroencephalogram (EEG)** recording and circuit manipulation to systematically analyze the correlation between this pathway and natural arousal, and distinguish its functional differences in the two physiological states.

In addition, although we supplemented experimental data from female mice in this study, we did not further explore the molecular mechanisms underlying gender differences. Sexual dimorphism widely exists in the nervous system, including the density of 5-HTergic neurons, receptor expression levels and synaptic transmission efficiency. These differences may lead to divergent responses of the CeA-mPFC circuit to anesthesia and interventions between male and female individuals. In follow-up research, we will adopt transcriptome sequencing to screen gender-specific regulatory factors (e.g., differentially expressed genes and non-coding RNAs) in the CeA-mPFC circuit, so as to provide a more comprehensive theoretical basis for personalized perioperative anesthesia management.

Regarding circuit interaction, the arousal process is coordinately regulated by multiple neural systems. The CeA-mPFC 5-HTergic circuit interacts extensively with other arousal-related networks, such as the LC noradrenergic system and basal forebrain cholinergic system. A recent study reported that the CeA-mPFC pathway may indirectly modulate arousal by regulating norepinephrine release. Specifically, CeA serotonergic neurons project to the LC and regulate the activity of noradrenergic neurons, thereby altering norepinephrine release in the mPFC and other downstream regions^65^. Meanwhile, cholinergic neurons in the basal forebrain form synaptic connections with the CeA and mPFC and may mutually regulate the function of the 5-HTergic circuit. Future studies will adopt multi-region simultaneous recording to detect the activity of CeA-mPFC 5-HTergic neurons and LC noradrenergic neurons, and clarify the synergistic or antagonistic relationships among different arousal circuits during anesthetic emergence.

This study also has several inherent limitations. First, consciousness recovery is modulated by the synergistic activity of multiple brain regions. The CeA and mPFC are important components of the ascending reticular activating system^43,44^, and the 5-HTergic circuit interacts with cholinergic arousal systems, GABAergic sleep-promoting systems, dopaminergic systems and noradrenergic systems, forming an extremely complex neural network^50ߝ52^. Second, 5-HT exerts bidirectional regulatory effects on sleep-wake activity, which depend on the activation state of the central serotonergic system, and 5-HT receptor subtypes are unevenly distributed across different brain regions^18,53ߝ55^. Third, 5-HT receptor subtypes possess complex functional characteristics^56^. Although blocking 5-HT₂A receptors with KET significantly delays anesthetic emergence, KET also weakly blocks α1-adrenergic receptors and histamine H1 receptors^20^. Accumulating evidence shows that 5-HT can activate GABAergic neurons via 5-HT₂A receptors to induce inhibitory postsynaptic currents^57^, indicating that the interaction between 5-HTergic and GABAergic systems during anesthetic emergence requires in-depth exploration. Moreover, other 5-HT receptor subtypes such as 5-HT₃R and 5-HT₆R have also been reported to promote arousal^58,59^. Comprehensive research is still needed to fully elucidate the neural mechanisms of consciousness transitions during general anesthesia.

We explicitly clarify that 5-HTP used in this study serves only as an experimental tool for activating the serotonergic pathway, rather than a candidate clinical therapeutic drug. Its core function is to verify the mechanistic link between enhanced 5-HT activity and accelerated anesthetic emergence, and it is not suitable for direct clinical application. As a systemic intervention, 5-HTP lacks tissue and cell-type specificity, so subsequent research needs to develop more targeted strategies for clinical transformation.

Furthermore, we discuss the potential limitations of all experimental tools used in this study and the corresponding quality control measures:1) Viral vector specificity: We acknowledge that the TPH2 promoter may cause low-level off-target expression. In this study, we combined pharmacological inhibition (e.g., p-chlorophenylalanine, PCPA, to block endogenous serotonin synthesis) and TPH2-Cre mouse models to exclude the interference of off-target cells, confirming that the observed physiological effects were specifically mediated by 5-HTergic neurons. 2) Chemogenetic tool limitation: CNO can be metabolized into clozapine in vivo and cause non-specific effects. Our pre-experiments verified that the CNO dose (1 mg/kg) used in this study did not produce obvious off-target activities such as activating dopamine D2 receptors, which is supported by published literature **(Gomez et al., 2017, *Science*)**. 3) Pharmacological antagonist selectivity: WAY-100635 and KET are not completely selective ligands. WAY-100635 has weak affinity for other 5-HT receptor subtypes, and KET exerts blocking effects on α1-adrenergic receptors. We fully considered these non-specific properties during data interpretation, and cross-validated all conclusions via optogenetics, chemogenetics and multiple pharmacological assays to guarantee result reliability.

To conclude, targeted activation of the CeA–mPFC serotonergic pathway via 5-HT₁A and 5-HT₂A receptors rescues the synaptic transmission deficits induced by anxiety and depression, and ultimately restores consciousness and accelerates sevoflurane emergence. This study not only deepens our understanding of neural mechanisms underlying consciousness transitions, but also validates the CeA-mPFC 5-HTergic circuit as a promising therapeutic target. It provides solid theoretical and experimental evidence for developing targeted interventions to improve anesthetic recovery in preoperative patients with anxiety and depression.

## 4 Materials and methods

### 4.1 Animals and PCR-based genotyping for Sert-Cre transgenic mice

This study was approved by the Animal Advisory Committee of Zhejiang University, and all experimental procedures adhered to the National Institutes of Health Guidelines for the Care and Use of Laboratory Animals. 8-week-old Wild-type C57BL/6 J mice were obtained from the Animal Experiment Center of Zhejiang University. The mice were housed and bred in the SPF-Class Housing of Laboratory of the Animal Center of Zhejiang University School of Medicine and maintained under standard conditions (indoor temperature 25°C, ambient humidity 65%, standard 12-h light/dark cycle) with ad libitum access to rodent food and water. To eliminate potential interference from gender and the female estrous cycle, only male mice were utilized, and all experiments were conducted between 9:00 and 15:00.

The generation of Sert-Cre transgenic mice with a pure C57BL/6J genetic background was achieved via successive intercrossing and genotyping screening. The original Sert-Cre transgenic founders maintained on the C57BL/6J background were used as the starting parental strain. Homozygous Sert-Cre mice were mated with wild-type C57BL/6J littermates to produce heterozygous F1 offspring carrying the Sert-Cre transgene. Serial breeding across multiple generations was performed, and all progeny were subjected to routine PCR genotyping after tissue sampling to distinguish homozygous Sert-Cre, heterozygous Sert-Cre, and wild-type mice. Heterozygous Sert-Cre mice obtained from successive intercrosses were selected for subsequent behavioral, fiber photometry, optogenetic and chemogenetic experiments.

To conduct genotyping, 3–5 mm toe biopsies were collected from each mouse at one week postnatal for genomic DNA extraction. Tissue fragments were transferred into sterile EP tubes, followed by DNA lysis with a mixed buffer system (AD1 buffer: AD2 buffer = 4: 1). Samples were incubated at room temperature for 10 min, then heated at 95 °C for 3 min to fully release genomic DNA, supplemented with AD3 buffer and mixed evenly. Prepared DNA lysates were stored at 4 °C and subjected to PCR amplification within 7 days.

The PCR reaction system contained 4 μl primer mixture, 12.5 μl Taq DNA polymerase master mix, 7 μl nuclease-free ddH₂O, and 1.5 μl tissue DNA template. Two distinct primer pairs were adopted for allele discrimination: sert-F + sert-R1 for wild-type alleles (predicted amplicon size: 295 bp), and sert-F + sert-R2 for mutant Sert-Cre transgenic alleles (predicted amplicon size: 195 bp). The primer sequences are listed below: sert-F: GAG CTC TCA GTC TTG TCT CCA, sert-R1: GAG TGT GGC GCT TCA TCC, sert-R2: AGG CAA ATT TGT TGT GCG. After PCR amplification, agarose gel electrophoresis was performed to separate target fragments. Briefly, 3 g agarose powder was dissolved in 100 mL 1×TAE buffer and heated in a microwave until fully dissolved. After cooling to approximately 60 °C, nucleic acid stain was added, and the gel solution was poured into casting molds with sample combs inserted. Once solidified, the gel was placed in an electrophoresis tank filled with TAE running buffer. PCR products were loaded into sample wells, and electrophoresis was run at a constant voltage of 170 V for 18 min. Post-electrophoresis gel imaging was performed to visualize DNA bands for genotype classification: wild-type mice: only a single 295 bp wild-type band (amplified by sert-F/sert-R1); heterozygous Sert-Cre mice: both 295 bp wild-type band and 195 bp mutant band; Homozygous Sert-Cre mice: only a single 195 bp transgenic mutant band (amplified by sert-F/sert-R2). Representative agarose gel genotyping results are provided in Supplementary Figure 2B.

### 4.2 Sevoflurane anesthesia induction and emergence time calculation

The mice were freely placed in an airtight inhalation anesthesia box (20 cm × 10 cm × 15 cm; V101, RWD Life Sciences, Inc., Shenzhen, China) for a 30-minute habituation period. The bottom of the box was preheated using an electric heating pad for 15 minutes prior to the experiment to ensure a consistent temperature throughout. Sevoflurane anesthesia was induced by administering 3% sevoflurane with 100% oxygen at a constant flow rate of 2 L/min. The righting reflex was evaluated by rotating the anesthesia box 90° every 15 seconds. LORR was considered when the mouse remained in a dorsally recumbent position and could not spontaneously return to an upright posture. Anesthesia was maintained for 30 minutes, after which sevoflurane administration was stopped, and rapid oxygenation (80% O_2_, 20% N_2_) at 2 L/min was used to remove residual sevoflurane from the box and pipeline. RORR was determined when the mouse could independently return to an upright posture with all four paws on the floor. The induction time was defined as the interval from the activation of the sevoflurane vaporizer to LORR. The emergence time was defined as the interval from the cessation of anesthesia to RORR. The same batch of mice was used for both experimental and control groups to minimize individual differences and environmental variability. Throughout the experiment, the mice were monitored closely to ensure that the environmental conditions and experimental parameters remained optimal for consistent and reproducible results.

### 4.3 Establishment and behavioral tests of model with anxiety- and depression-like symptoms

**CRS model:** Mice were placed daily in a plastic air-accessible cylinder for 6 h for 21 consecutive days to establish anxiety model^60^. The size of the cylinder was similar to that of the animal, which made the animal almost immobile in the cylinder. The non-stressed controls were moved from the home cage to a test room without access food and water for 6 h. **LPS-induced model:** Acute treatment with the cytokine inducer LPS is also widely accepted in anxiety models^61^. LPS from Escherichia coli (serotype 0111: B4) was obtained from Sigma-Aldrich (St. Louis, MO, USA). For the experiments, LPS was administered intraperitoneally at a dose of 1 mg/kg. The LPS was dissolved in saline (0.9% NaCl), while the control group received an IP injection of saline at a volume of 10 ml/kg. **Open field test (OFT):** The open field chamber, measuring 50 cm × 50 cm, was constructed from transparent plastic with a 25 cm × 25 cm centrally marked square^62^. Each mouse was placed in the center of the chamber, and their behavior was recorded over a 10 min period using ANY Maze video tracking system (SD Instruments, CA, USA). The apparatus was cleaned with 75% alcohol after the test. During the experiment, the time spent in the center area, and total distance traveled were continuously monitored and analyzed. **Elevated plus maze (EPM):** The EPM test is a common method for assessing anxiety-like behavior in mice^63,64^. The maze consists of two opposite open arms (30 cm × 5 cm) and two enclosed arms (30 cm × 5 cm) extending from a central platform (5 cm × 5 cm), positioned 74 cm above the floor. During the test, each mouse is placed in the center of the maze, facing an open arm, and observed for 5 min. The number of entries into the open arms and the time spent there were recorded by ANY Maze video tracking system. After each trial, the apparatus is cleaned with 75% ethanol. **Tail suspension test (TST):** The TST apparatus consisted of a hook positioned 60 cm above the ground. Mice were suspended from the hook by their tail using adhesive tape. Immobility was measured during the final 4 minutes of a 6-minute test session^63^. **Forced swim test (FST):** The FST apparatus consisted of a transparent cylinder (12 cm diameter, 25 cm height) containing water (22°C-24°C). The water depth was set to prevent the mice from touching the bottom with their tails or hind limbs. The mice were placed in the water and their behavior was recorded for 6 minutes, with immobility analyzed in the last 4 minutes^63^.

### 4.4 Stereotaxic surgery and virus microinjection surgery

8-week-old C57BL/6J mice were weighed and anesthetized with 1% pentobarbital sodium (50 mg/kg) and fixed in a stereotaxic apparatus (68018, RWD Life Sciences, Shenzhen, China). The body temperature of the anesthetized mice was maintained at 37°C using a heating pad and an ophthalmic ointment was applied to their eyes to avoid dryness. Carefully expose the skull by making a midline incision, then remove dirt using 3% hydrogen peroxide, and finally rinse away residue with saline solution. To ensure precise positioning, the skull was leveled using a microscope, and the coordinates were confirmed based on the fourth edition of the Mouse Brain Atlas (Paxinos and Franklin, 2013). Virus injection was performed at the following coordinates for specific brain regions: for the CeA (AP: −0.37 mm, ML: ±2.80 mm, DV: −4.95 mm), for the mPFC (AP: +0.90 mm, ML: ±0.45 mm, DV: −1.50 mm) and for the lateral ventricular (AP: −0.47 mm, ML: −1.00 mm, DV: −2.40 mm). A miniature cranial drill was used to create a small hole in the skull, avoiding damage to underlying tissues. Virus was injected at a rate of 20-40 nL/min using an Ultra Micro Pump (160494 F10E, WPI). A total of 100 nL virus was delivered to each site, and the needle remained in place for 10 minutes post-injection to allow for virus diffusion. After the injection, the needle was withdrawn slowly, and the incision was sutured. According to the experimental needs, optical fibers or cannulas were subsequently implanted in the corresponding location and secured with dental cement. After that, the incision was sutured, and the mice were placed on a heating blanket to be transferred to a cage when the mice were awake. To allow for virus expression, animals were kept for 3 weeks before further experimentation. At the end of the experiment, viral expression and the location of optical fibers or cannulas implantation were confirmed by immunohistochemistry.

### 4.5 Retrograde tracing

C57BL/6Jmice at 8 weeks of age were anesthetized with 1% pentobarbital sodium (50 mg/kg) and fixed the head with a stereotactic apparatus (68018, RWD Life Sciences, Inc., Shenzhen, China), as previously described. As for the retrograde tracing from the CeA to the mPFC: 100nl of Cholera Toxin Subunit B-555 (CTB-555) (1µg/µL, BrainVTATechnology Co., Ltd., Wuhan, China) was injected in the mPFC with a total content of 100 nL for retrograde labeling of projection neurons. The following experiments were performed at least one week after surgery to ensure the expression of virus. As for the RV virus cross-synaptic tracing from the mPFC to the CeA: the TPH2-dependent helper viral vectors (rAAV-EF1a-DIO-mCherry-F2A-TVA-WPRE-hGE polyA, rAAV-EF1a-DIO-oRVG -WPRE-he polyA and rAAV-TPH2-CRE-WPRE-hGE polyA) (100nL, titer ≥ 5^12^ vg/mL, Brain VTA Technology Co., Ltd., Wuhan, China) were injected into the CeA. Three weeks later, RV-EnvA-AG-EGFP (100nL, titer: 2^8^ IFU/mL, Brain VTA Technology Co., Ltd., Wuhan, China) was injected into the CeA. Seven days later, sacrificed the mice and observed RV-labeled neurons in the mPFC.

### 4.6 Optogenetics

Three weeks before the experiment began, the CeA or mPFC was microinjected with an optogenetic virus of 100 nL of pAAV-TPH2 PRO-ChETA-EYFP-WPRES-PAS (100 nL, titer: 10^13^ vg/mL, Shanghai Sunbio Medical Biotechnology Co., Ltd.). One week before the experiment began, the CeA or mPFC was implanted with optical fibers (FOC-W-1.25-200-0.37-3.0, Yingbo Technology Co., Ltd., Hangzhou, China). TPH2-ChETA expression in the CeA or mPFC was determined by immunohistochemistry after the completion of optogenetic experiments. The mice were placed into the anesthesia box for 30 min before anesthesia induction, and 20 mins later, they were exposed to blue light photostimulation (465 nm, 20 Hz, 20 ms pulse, 5 mW / 10 mW / 15 mW / 20mW) for 10 minutes. We finally identified 15 mW to activate CeA^5-HT^ neurons and 20mW to activate mPFC^5-HT^ neurons as the optimal optogenetic activation parameter and used it in the subsequent experiments.

### 4.7 Chemogenetics

Three weeks before starting the experiment, chemogenetic viruses of 100 nL of rAAV-TPH2-hM3D(Gq)-EGFP-WPREs or rAAV-TPH2-hM4D(Gi)-mCherry-WPREs (100 nL, titer: 10^12^ vg/mL, Brain VTA Technology Co, Ltd) were microinjected into the CeA or mPFC, and TPH2-hM3D or TPH2-hM4D expression in the CeA or mPFC was determined by immunohistochemistry after the completion of chemogenetic experiments. Clozapine-N-oxide (CNO) (MedChemExpress, HY-17366) were dissolved in saline and injected intraperitoneally 10 minutes before sevoflurane anesthesia to activate neurons using the chemogenetics approach. In this study, we tested different doses of CNO: 0.25 / 0.5 / 1.0 / 1.5 / 2.0 mg/kg. We concluded that 0.5 mg/kg to activate CeA^5-HT^ neurons and 1.0 mg/kg to activate mPFC^5-HT^ neurons in normal mice, 1.0 mg/kg to activate CeA^5-HT^ neurons and 2.0 mg/kg to activate mPFC^5-HT^ neurons in CRS and LPS mice was the optimal dose and used this dose for the subsequent IP administration.

### 4.8 Fiber photometry

For the photometry recordings, rAAV-TPH2-GCaMP6s-WPRE-hGH-pA virus (100nL, titer: 10^12^vg/mL, Brain VTA Technology Co., Ltd., Wuhan, China) was injected into the CeA or mPFC. Two weeks later, the optical fiber (FOC-W-1.25-200-0.37-4.0, Inper, Hangzhou, China) was implanted in the same area located 0.05 mm above the virus injection point (AP: −0.37 mm, ML: ±2.80 mm, DV: −4.90 mm/AP: +0.90 mm, ML: ±0.45 mm, DV: −1.45 mm) and fixed in place with dental cement. Then, 1 week after fiber optic placement, the calcium activity of the target neurons was monitored. Fluorescence emissions were recorded using a fiber photometry system (Inper, C11946) using a 488 nm diode laser, and the mice were placed in a dark box to avoid the influence of natural light source of calcium signal recording and accessed to the computer recording software to record calcium signals. After the recording was completed, the results recorded by the optical fiber were read into the calcium signaling data analysis software. For the experiment to investigate changes in calcium signaling during different stages of sevoflurane anesthesia, we recorded calcium signaling changes in a total of six mice and finally selected representative data. In this part, we divided the whole calcium signal recording process into four stages: Wakefulness, Before LORR, Sevoflurane, and Emergence. The time points for awake, turn on the sevoflurane vaporizer, LORR and turn off the sevoflurane vaporizer in mice were respectively selected as time = 0. The selected traces lasting for 16 s were based on the length of a complete Ca^2+^ signal. The calcium signal intensity was calculated as the value of ΔF/F = (F-F0)/F0.

### 4.9 Pharmacological experiment

C57BL/6 J mice were intraperitoneally injected with 5-HTP (25 / 50 / 75 / 100 / 125 mg/kg) and re-injected 24 h later. One hour after the second injection, mice were placed into the anesthesia box for further experiments. The ICV injection, intra-CeA or intra-mPFC 5-HTP microinjection experiments were conducted one week after the implantation of cannulas. For implantation of cannula in lateral ventricle and nucleus, the lateral ventricle cannula (O.D.0.30mm/C.C2.4/M3.5, 62004, RWD Life Science, China) and CeA cannula (O.D.0.50mm/C.C2.4/M3.5, 62004, RWD Life Science, China) were employed. Secure the cannula, screws, and skull tightly with dental cement and powder. The coordinates of the target brain regions involved in this experiment were CeA (AP: −0.37mm, ML: ±2.80mm, DV:−4.95mm), mPFC (AP: +0.90mm, ML: ±0.45mm, DV: −1.50mm) and the lateral ventricular (AP: −0.47 mm, ML: −1.00 mm, DV: −2.40 mm). In the experiments, the volume of saline or drug injected into the lateral ventricle was 2 μL, and the volume of saline or drug microinjected into the cannula was 200 nL. 5-HTP was injected into the lateral ventricular (11.35 / 22.7 mol/L) and the CeA or the mPFC cannulas (2.5 / 5.0 / 1.0 mg/mL), and sevoflurane anesthesia was initiated 15 minutes later. According to the experimental needs, one week after cannulas implantation, the mice were injected with 5HT1A receptor (5-HT1AR) antagonist WAY-100635 (WAY), 5-HT2A receptor (5-HT2AR) antagonist KET through the lateral ventricle, CeA, and mPFC cannulas.

### 4.10 Immunohistochemistry

For immunohistochemistry analysis of the brain, the mice were anesthetized and transcranial perfuse with phosphate-buffered saline (PBS) followed by a 4 % paraformaldehyde solution. The brains were carefully extracted, post-fixed in 4% paraformaldehyde for an additional 6 h at 4°C, and then dehydrated in 30% sucrose solution for 24 h at 4°C. Subsequently, the brain was frozen in the Optimal Cutting Temperature compound and sectioned into 35-μm coronal slices using a cryostat microtome (CM30503, Leica Biosystems, Buffalo Grove, IL, USA). Frozen slices were washed three times in PBS for 5 min and incubated in a blocking solution containing 10% normal donkey serum (017-000-121, Jackson ImmunoResearch, West Grove, PA), 1% bovine serum albumin (A2153, Sigma-Aldrich), and 0.3% Triton X-100 in PBS for 2 h at room temperature. The sections were incubated with primary antibodies at 4°C overnight; this was followed by incubation in a solution of secondary antibodies for 2 h at room temperature. The primary antibodies used were Rabbit anti-TPH2 (1:200, 100-74555, Novus Biologicals), Mouse anti-TPH2 (1:100, T0678, Sigma-Aldrich), and Rabbit anti-c-Fos (1:1000, 2250T, Cell Signaling Technology), and the secondary antibodies used were Donkey anti-Rabbit Alexa Fluor Cy5 (1:1000; A10523, Thermo Fisher Scientific), and Donkey anti-Mouse Alexa Fluor 546 (1:1000; A10036, Thermo Fisher Scientific). The brain slices were washed three times with PBS for 15 min, and then they were deposited on glass slides and incubated in a DAPI solution at room temperature for 7 min. Finally, an anti-fluorescence attenuating tablet was applied to seal the slides. Confocal images were acquired using a Nikon A1 laser-scanning confocal microscope (Nikon, Japan) or a VS120 microscope (Olympus, Japan), and further image processing was done using Image J (NIH, Baltimore).

### 4.11 Quantification and statistical analysis

All the experimental data were reported as mean ± SEM. The Shapiro-Wilk normality test was performed for all experimental data before data analysis. For comparative analyses of the two groups, normally distributed data were compared using the Student’s T-test, including independent samples T-test and paired samples T-test, and non-normally distributed data were compared using the Mann-Whitney U test or Wilcoxon signed-rank test. The Levene test was used to detect the homogeneity of square difference. After the data met the normal distribution and homogeneity of variance, ordinary one-way ANOVA analysis was used for comparing three or more groups, and two-way ANOVA analysis followed by Bonferroni’s test were used for comparisons between two factors and different groups. Statistical significance was indicated by *P* < 0.05. GraphPad Prism 8.0 (GraphPad Software, Inc) and SPSS 26.0 (SPSS, Inc) were used for data visualization and statistical analysis, respectively.

All behavioral test data were analyzed using GraphPad Prism 9.0 software. For multiple group comparisons, one-way or two-way analysis of variance (ANOVA) was performed, followed by Tukey’s post hoc test or Bonferroni correction to eliminate type I errors. All quantitative data are presented as mean ± standard error of the mean (SEM). A P value < 0.05 was considered statistically significant. All corrected statistical markers have been updated in all relevant figures and legends.

## Authors’ contributions

H.H.Z. and X.Z.C. Conceptualization, Resources, Data curation, Supervision, Funding acquisition, Validation, Project administration, Writing – review and editing; Y.L.W., Conceptualization, Investigation, Writing – review and editing; Y.Y., Y.X.W., X.X.X., Data curation, Investigation, Writing – original draft; W.H.S., S.L. Software, Investigation, Writing – original draft; Q.X., L.Y.G., L.L., Z.W.Z., X.Y.D., Investigation, Writing – original draft; H.W, D.D.Y and Y.S., Methodology, Writing – original draft.

## Acknowledgments

We thank YuDong Zhou and Yi Shen for their help in experimental design.

## Data Availability

The data supporting the findings of this study are available within the article. Data will be made available upon reasonable request.

## Declarations

### Ethics Approval

All procedures were in accordance with the National Institutes of Health Guidelines for the Care and Use of Laboratory Animals and were approved by the Animal Advisory Committee of Zhejiang University.

### Consent to Participate

Not applicable.

### Consent for Publication

Not applicable.

## Supplementary information

**Supplementary Figure 1.**
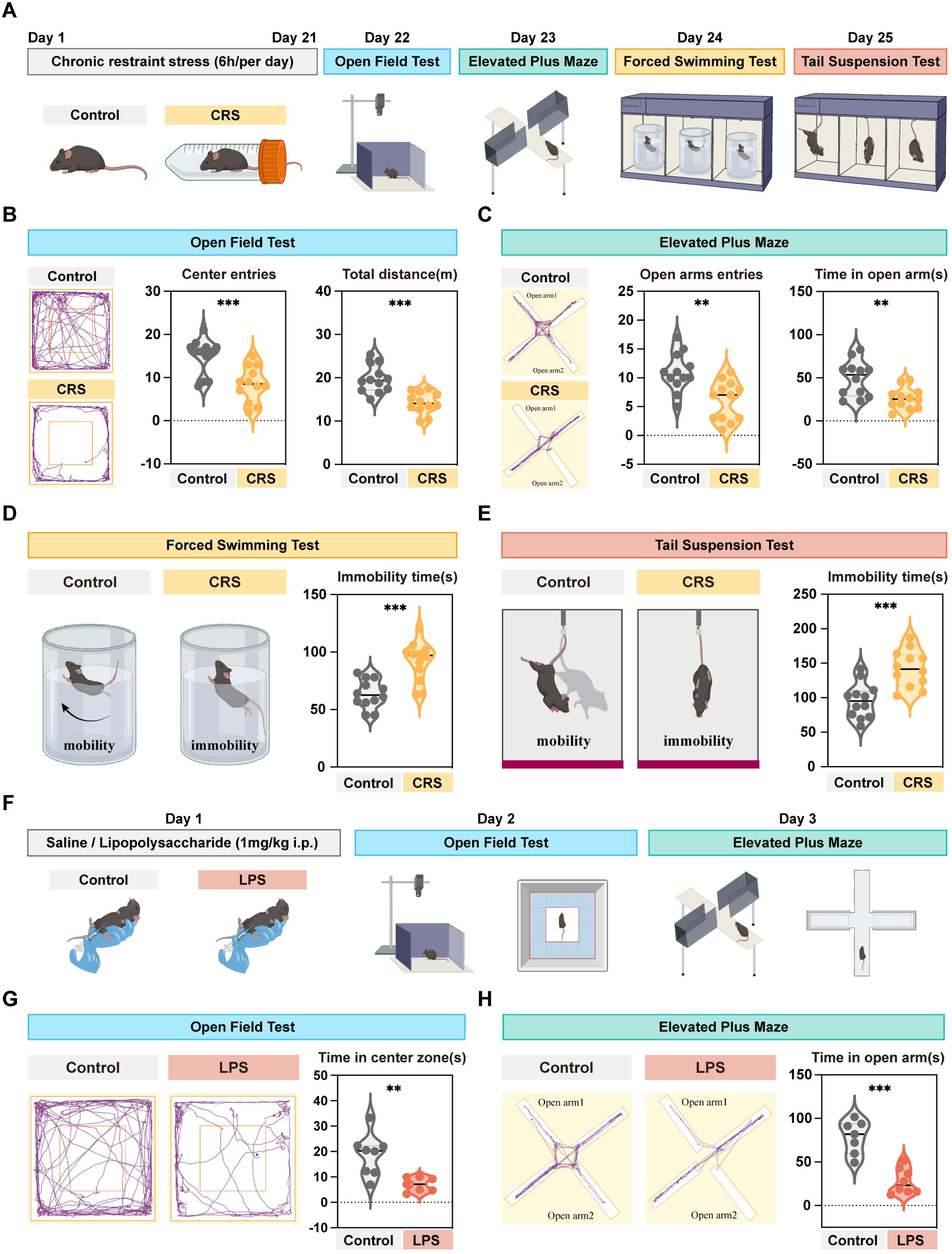
Establishment of CRS and LPS-induced model and the assessment of behavioral tests for anxiety- and depression-like symptoms. **(A)** Schematic of the experimental protocol for the establishment of CRS model and behavioral assessment of anxiety- and depression-like symptoms. **(B)** Representative locomotion tracks in OFT for the Control and CRS groups. Statistical analysis of the number of entries into the center field, and the total distance of the OFT for the Control and CRS groups. **(C)** Representative locomotion tracks in EPM for the Control and CRS groups. Statistical analysis of the time spent in and the number of entries to the open arm in the EPM for the Control and CRS groups. **(D)** Representative schematic of the FST for the Control and CRS groups. Statistical analysis of the duration of immobility in the FST for the Control and CRS groups. **(E)** Representative tracking of activity in the TST for the Control and CRS groups. Statistical analysis of the duration of immobility in the TST for the Control and CRS groups. **(F)** Schematic of the experimental protocol for the establishment of LPS model and behavioral testing assessment of anxiety- and depression-like symptoms. **(G)** Representative locomotion tracks in OFT for the Control and LPS groups. Statistical analysis of the time spent in the center zone of the OFT for the Control and LPS groups. **(H)** Representative locomotion tracks in EPM for the Control and LPS groups. Statistical analysis of the time spent in the open arm of the EPM for the Control and LPS groups. ***p < 0.001, **p < 0.01; Data are mean ± SEM; CRS, chronic restraint stress; LPS, lipopolysaccharide; OFT, open field test; EPM, elevated plus maze; FST, forced swimming test; TST, tail suspension test

**Supplementary Figure 2.**
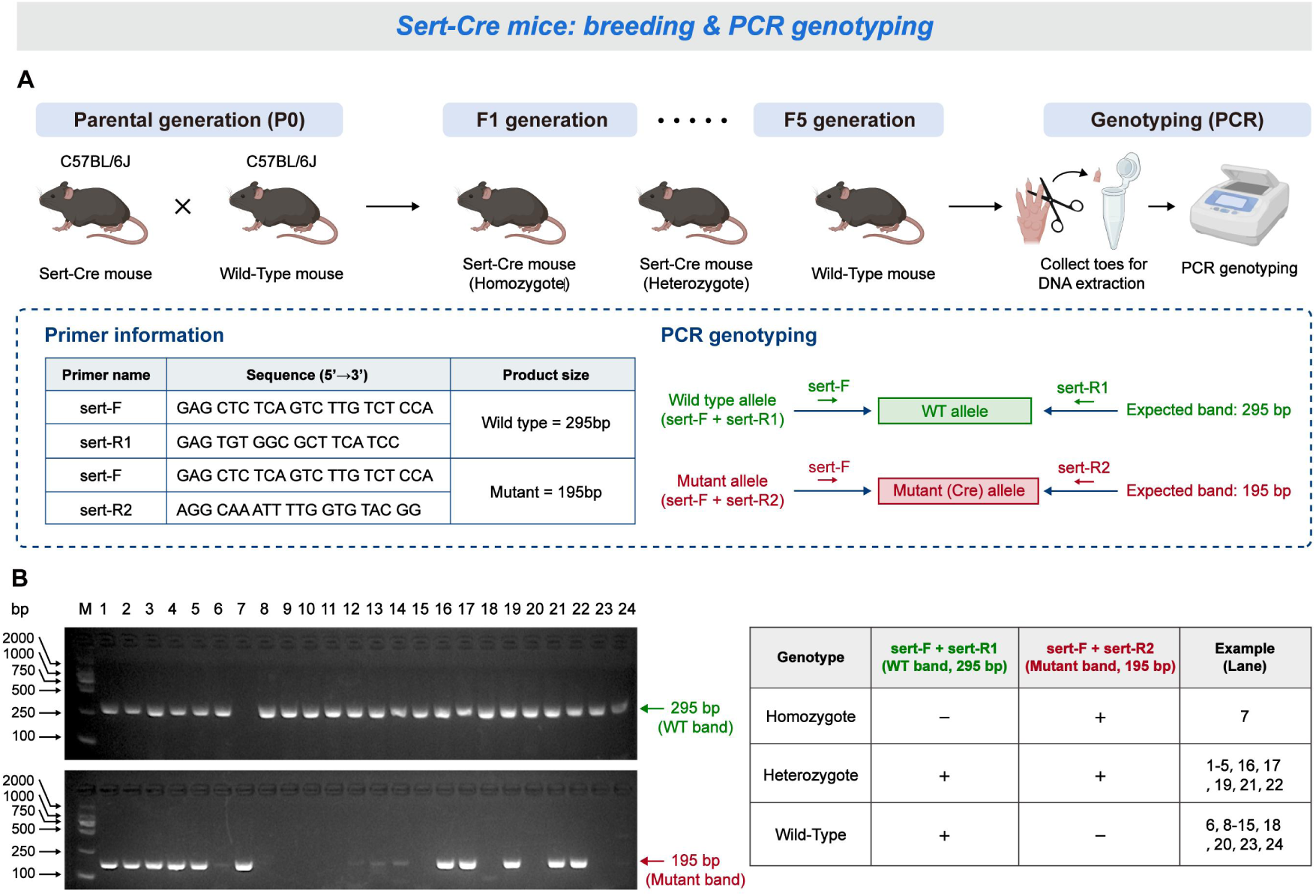
Breeding strategy and PCR genotyping of Sert-Cre mice. **(A)** Schematic illustration of the breeding strategy and PCR genotyping procedure for Sert-Cre mice. Sert-Cre mice on a C57BL/6J background were crossed with wild-type C57BL/6J mice to obtain offspring with different genotypes. Toe tissue samples were collected for DNA extraction, followed by PCR genotyping. The wild-type allele was amplified using sert-F and sert-R1, resulting in an expected band of 295 bp, whereas the mutant allele was amplified using sert-F and sert-R2, resulting in an expected band of 195 bp. **(B)** Representative agarose gel electrophoresis images showing PCR genotyping results from Sert-Cre mice. The upper gel shows the wild-type band of 295 bp, and the lower gel shows the mutant band of 195 bp. Genotypes were identified according to the presence or absence of these two bands: homozygous mice showed only the mutant band, heterozygous mice showed both wild-type and mutant bands, and wild-type mice showed only the wild-type band. Representative lane numbers for each genotype are summarized in the table. M, DNA marker; WT, wild-type; Sert, serotonin transporter

**Supplementary Figure 3.**
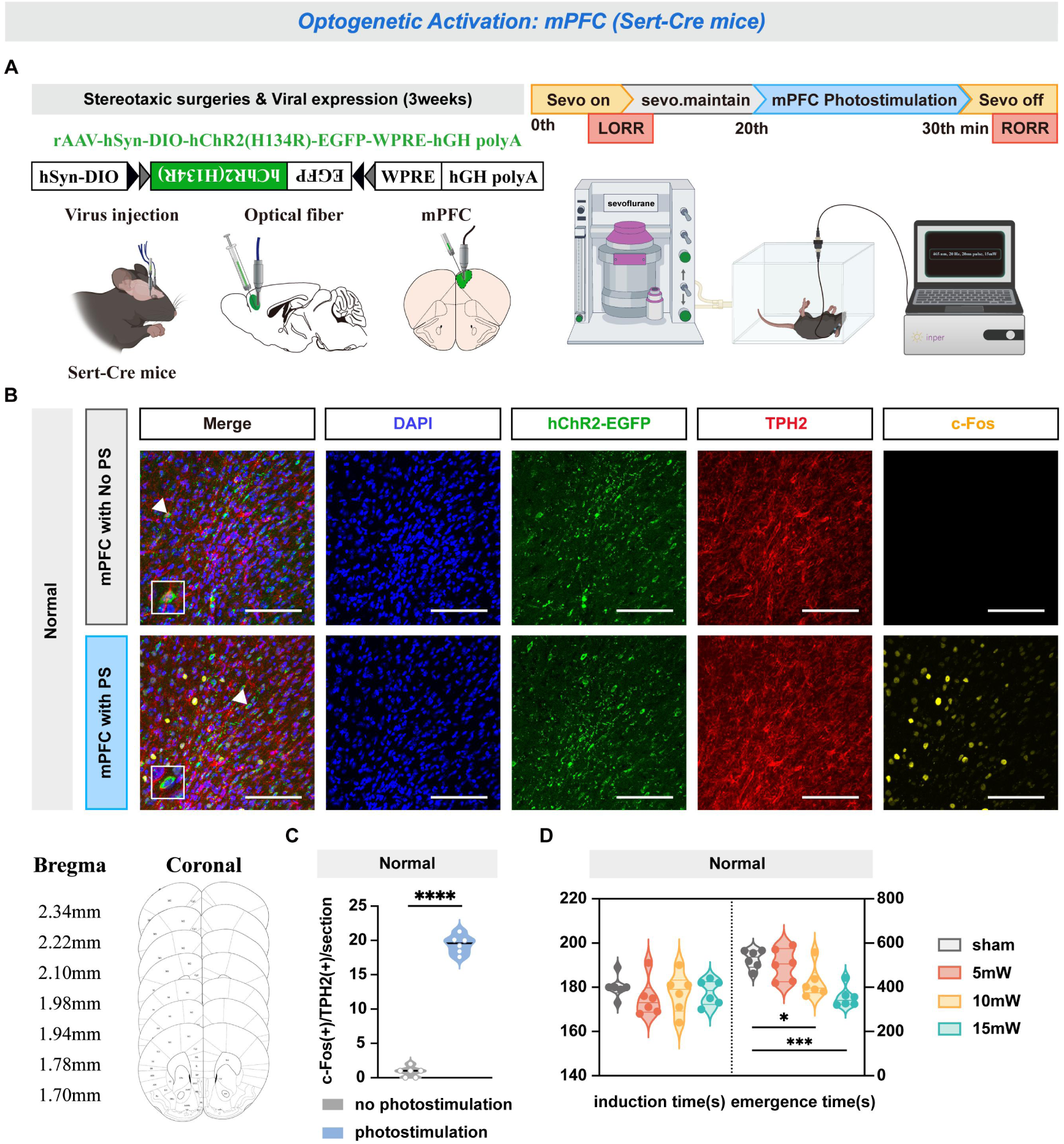
Optogenetic activation of mPFC^5-HT^ neurons promotes arousal from sevoflurane anesthesia in normal Sert-Cre mice. **(A)** Experimental protocol for optogenetic activation of mPFC^5-HT^ neurons in normal Sert-Cre mice. The schematic shows the position of the virus (rAAV-hSyn-DIO-hChR2(H134R)-EGFP-WPRE-hGH polyA) injection and optic fiber implantation in the mPFC. **(B)** Representative images of staining for c-Fos, TPH2, hChR2-EGFP and DAPI in the mPFC of the normal Sert-Cre mice with and without mPFC photostimulation. **(C)** The quantification of c-Fos(+)/TPH2(+)/section in the mPFC with or without mPFC photostimulation. **(D)** Induction time and emergence time for normal Sert-Cre mice in the sham, 5 mW, 10 mW and 15 mW groups. ****p < 0.0001; ***p < 0.001; *p < 0.05; Scale bar: 100 μm; Data are mean ± SEM; mPFC, medial prefrontal cortex; Sevo, sevoflurane; LORR, loss of righting reflex; RORR, recovery of righting reflex; Sert, serotonin transporter

**Supplementary Figure 4.**
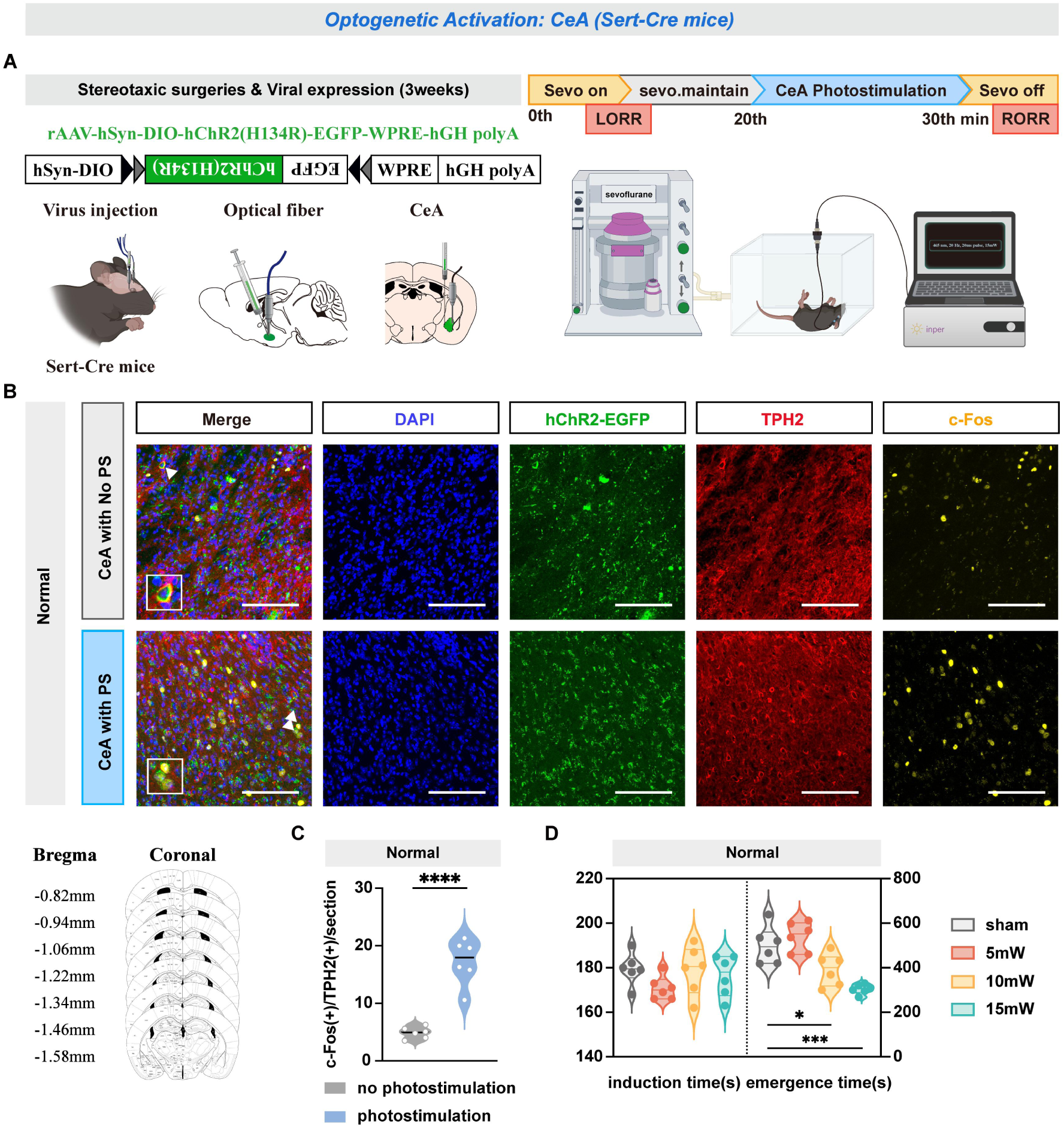
Optogenetic activation of CeA^5-HT^ neurons promotes arousal from sevoflurane anesthesia in normal Sert-Cre mice. **(A)** Experimental protocol for optogenetic activation of CeA^5-HT^ neurons in normal Sert-Cre mice. The schematic shows the position of the virus (rAAV-hSyn-DIO-hChR2(H134R)-EGFP-WPRE-hGH polyA) injection and optic fiber implantation in the CeA. **(B)** Representative images of staining for c-Fos, TPH2, hChR2-EGFP and DAPI in the CeA of the normal Sert-Cre mice with and without CeA photostimulation. **(C)** The quantification of c-Fos(+)/TPH2(+)/section in the CeA with or without CeA photostimulation. **(D)** Induction time and emergence time for normal Sert-Cre mice in the sham, 5 mW, 10 mW and 15 mW groups. ****p < 0.0001; ***p < 0.001; *p < 0.05; Scale bar: 100 μm; Data are mean ± SEM; CeA, central amygdala nucleus; Sevo, sevoflurane; LORR, loss of righting reflex; RORR, recovery of righting reflex; Sert, serotonin transporter

**Supplementary Figure 5.**
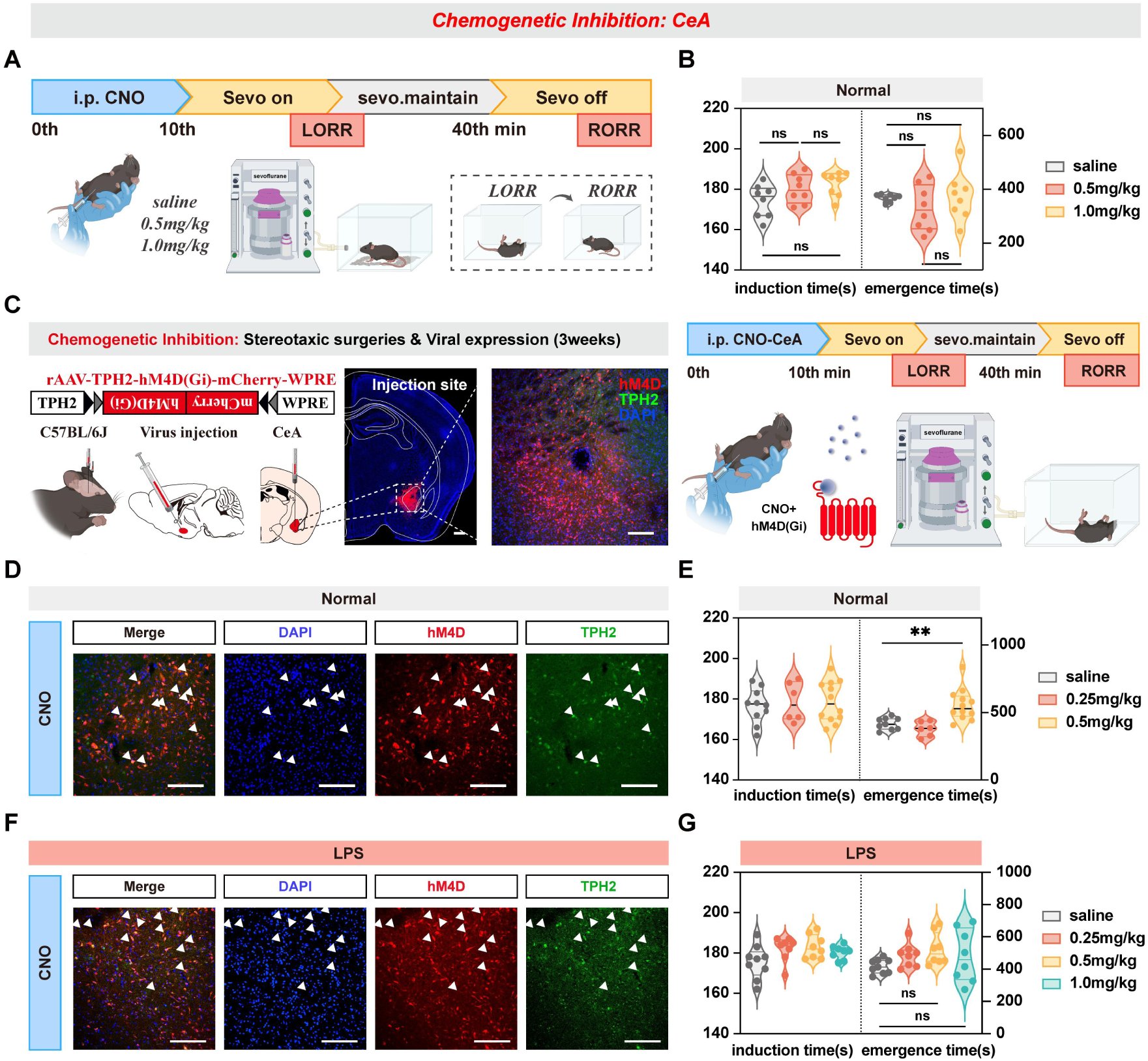
Chemogenetic inhibition of CeA^5-HT^ neurons prolonged the emergence time in normal mice while there is no effect on the emergence time in LPS mice. **(A)** Experimental protocol for i.p. injection of different doses of CNO. **(B)** Induction time and emergence time for i.p. injection in the saline control, 0.5 mg/kg CNO and 1.0 mg/kg CNO groups. **(C)** Experimental protocol for chemogenetic inhibition of CeA^5-HT^ neurons. Schematic of the position of the virus injection in the CeA, and representative coronal brain slice, showing the location of virus (rAAV-TPH2-hM4D(Gi)-mCherry-WPREs) injected in the CeA. **(D)** Representative images show the co-expression of and staining for hM4D, TPH2 and DAPI in the CeA of normal mice with CNO activation. **(E)** Induction time and emergence time for normal mice for i.p. injection in the saline control, 0.25 mg/kg CNO, 0.50 mg/kg CNO groups. **(F)** Representative images show the co-expression of and staining for hM4D, TPH2 and DAPI in the CeA of LPS mice with CNO activation. **(G)** Induction time and emergence time for LPS mice for i.p. injection in the saline control, 0.25 mg/kg CNO, 0.50 mg/kg and 1.0 mg/kg CNO groups. ns, p > 0.05, **p < 0.01; Scale bar: 100 μm; Data are mean ± SEM; i.p., intraperitoneal injection; CNO, clozapine-N-oxide; CeA, central amygdala nucleus; Sevo, sevoflurane; LORR, loss of righting reflex; RORR, recovery of righting reflex; LPS, lipopolysaccharide

**Supplementary Figure 6.**
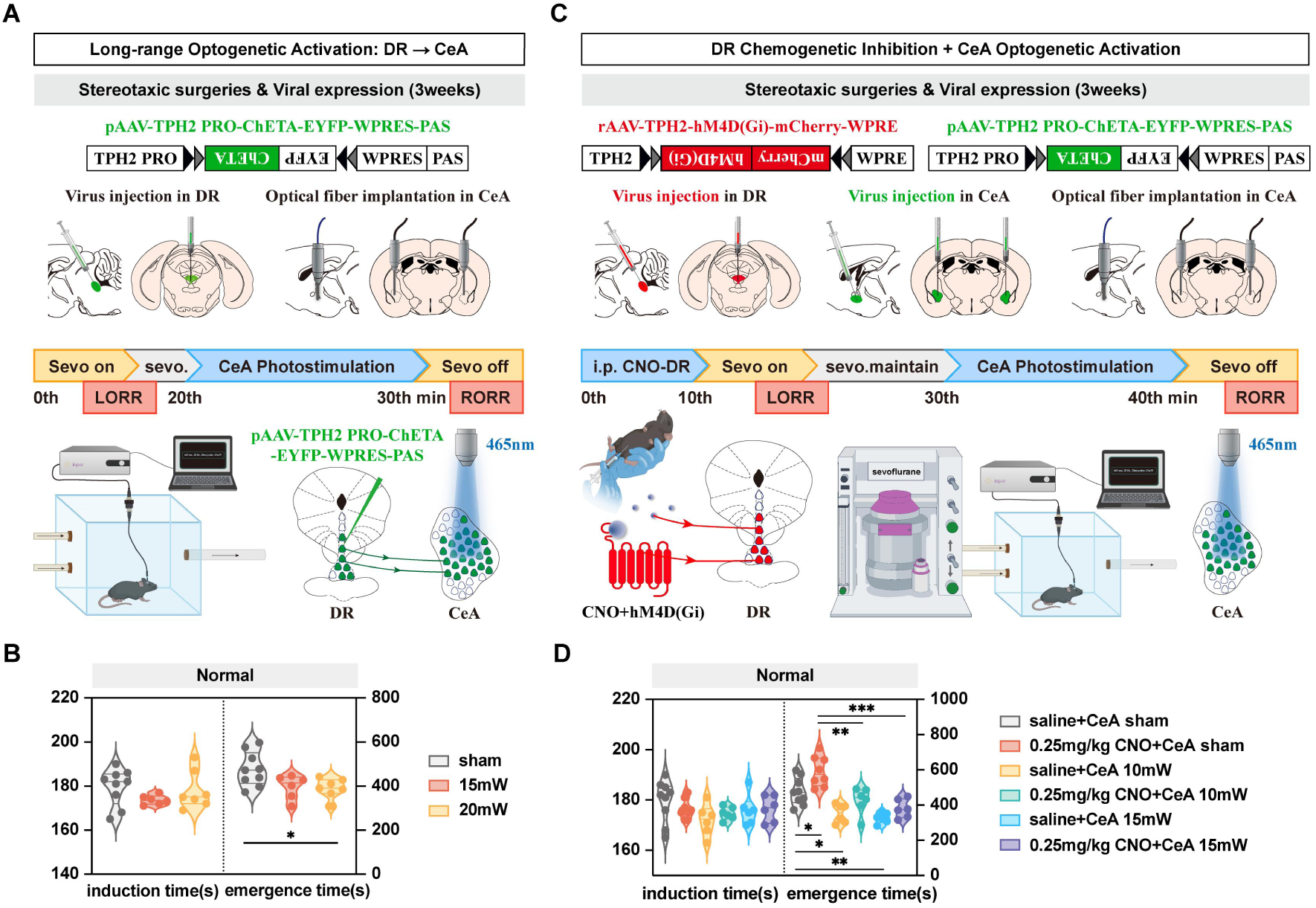
The pro-arousal effects of optogenetic activation of CeA^5-HT^ neurons regardless of the activity of DR^5-HT^ neurons. **(A)** Experimental protocol for optogenetic activation of the 5-HTergic nerve fibers projecting from the DR to the CeA. Schematic of the position of the virus (pAAV-TPH2 PRO-ChETA-EYFP-WPRES-PAS) injection in the DR and the optical fiber implantation in the CeA. **(B)** Induction time and emergence time for normal mice in the sham, 15 mW and 20 mW groups. **(C)** Experimental protocol for optogenetic activation of CeA^5-HT^ neurons with chemogenetic inhibition of the 5-HTergic nerve fibers projecting from the DR to the CeA. Schematic of the position of the virus (pAAV-TPH2 PRO-ChETA-EYFP-WPRES-PAS; rAAV-TPH2-hM4D(Gi)-mCherry-WPREs) injection and the optical fiber implantation. **(D)** Induction time and emergence time for normal mice for the saline + CeA sham, 0.25 mg/kg CNO + CeA sham, saline + CeA 10 mW, 0.25 mg/kg CNO + CeA 10 mW, saline + CeA 15 mW and 0.25 mg/kg CNO + CeA 15 mW groups. ***p < 0.001, **p < 0.01, *p < 0.05; Data are mean ± SEM; i.p., intraperitoneal injection; CNO, clozapine-N-oxide; CeA, central amygdala nucleus; DR, dorsal raphe nucleus; Sevo, sevoflurane; LORR, loss of righting reflex; RORR, recovery of righting reflex

**Supplementary Table 1.**
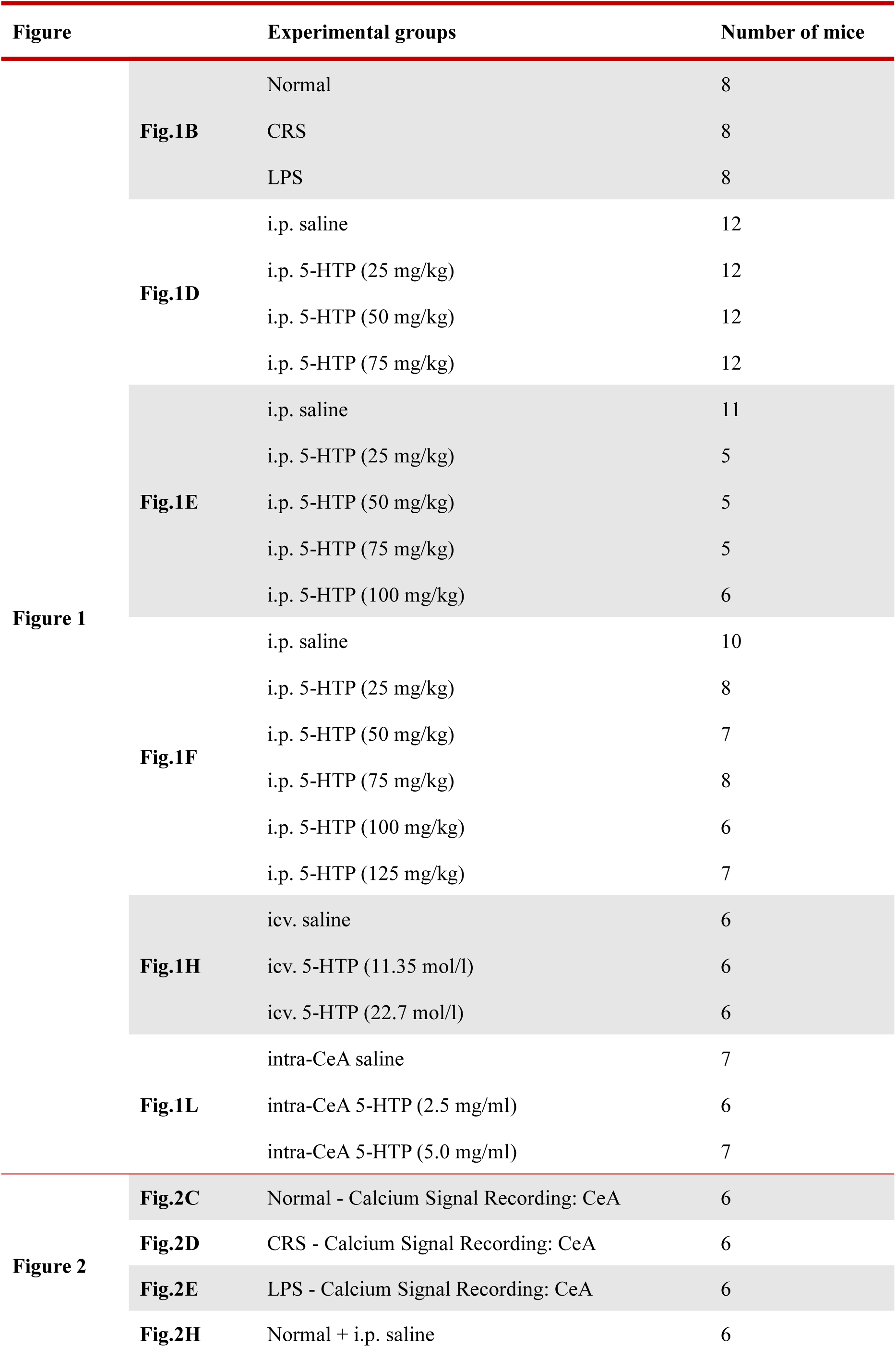
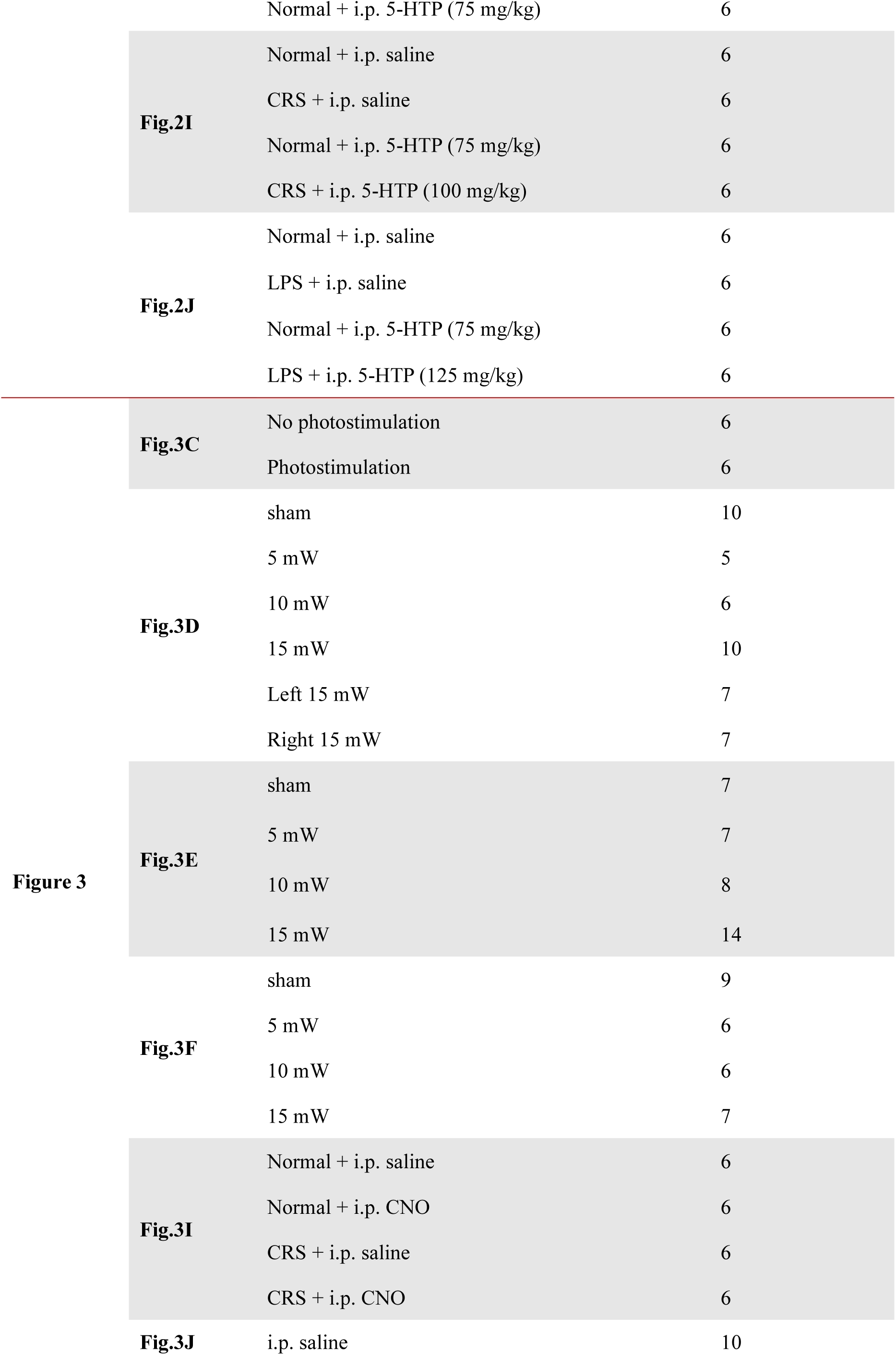
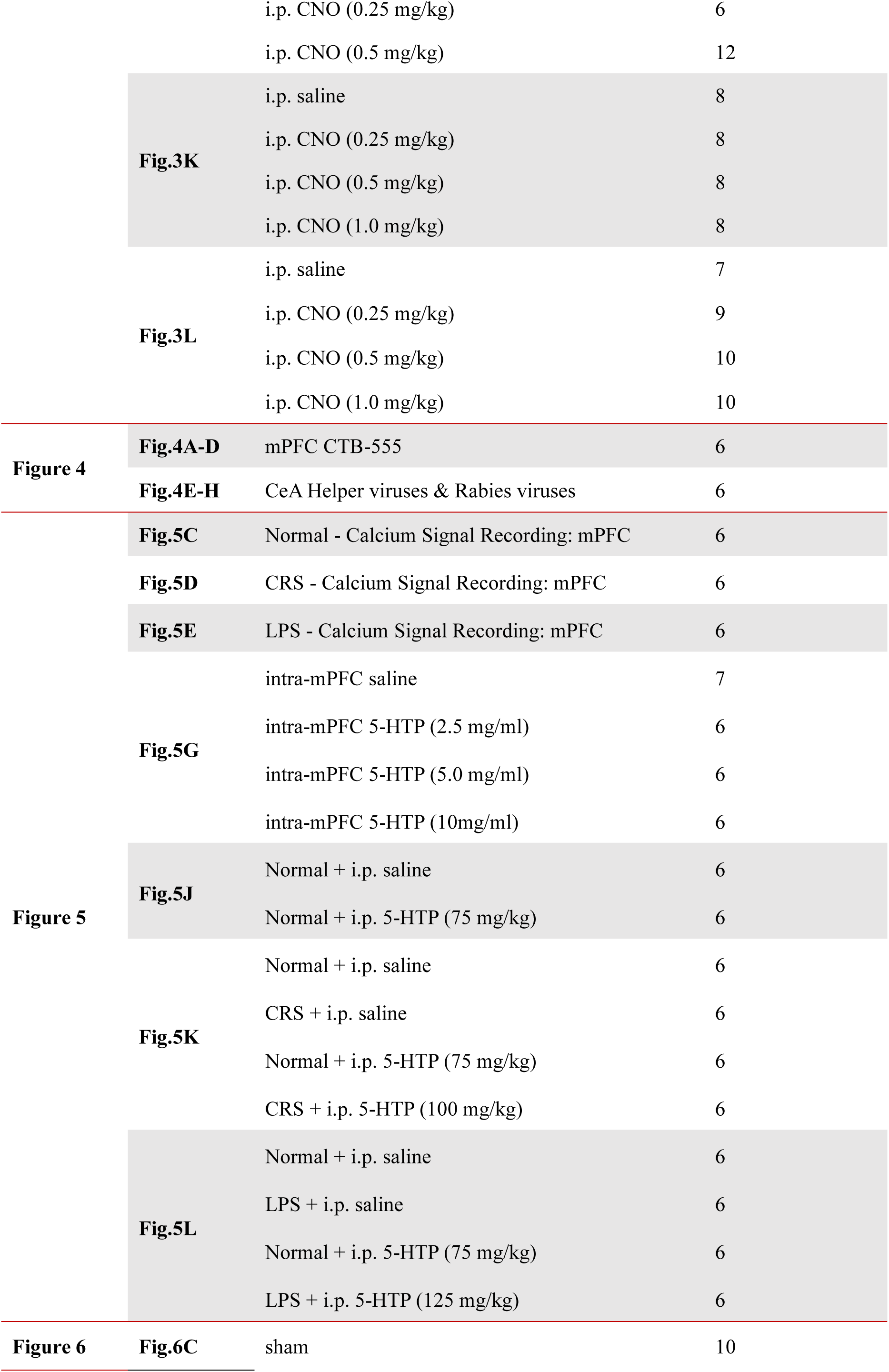
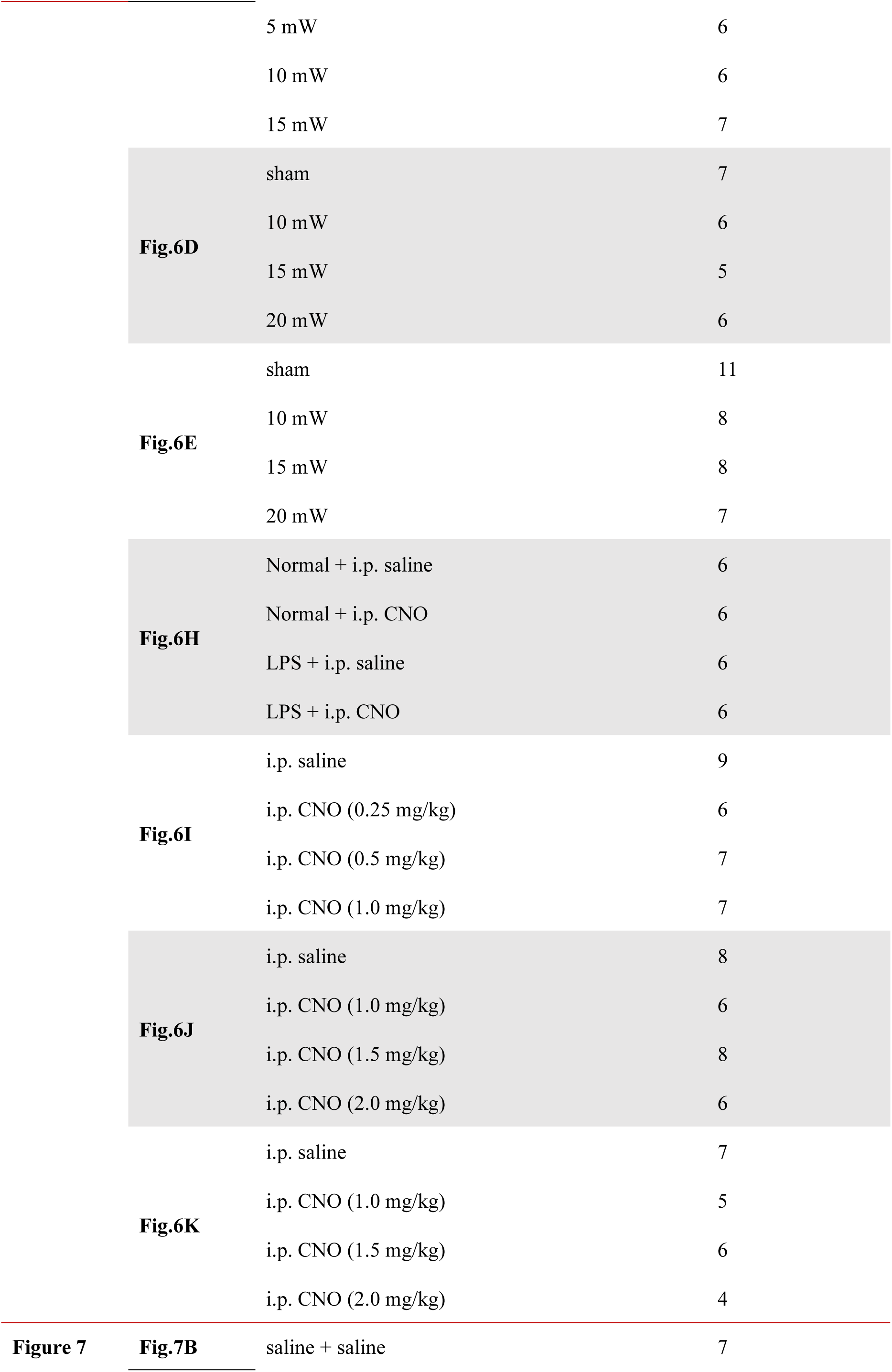
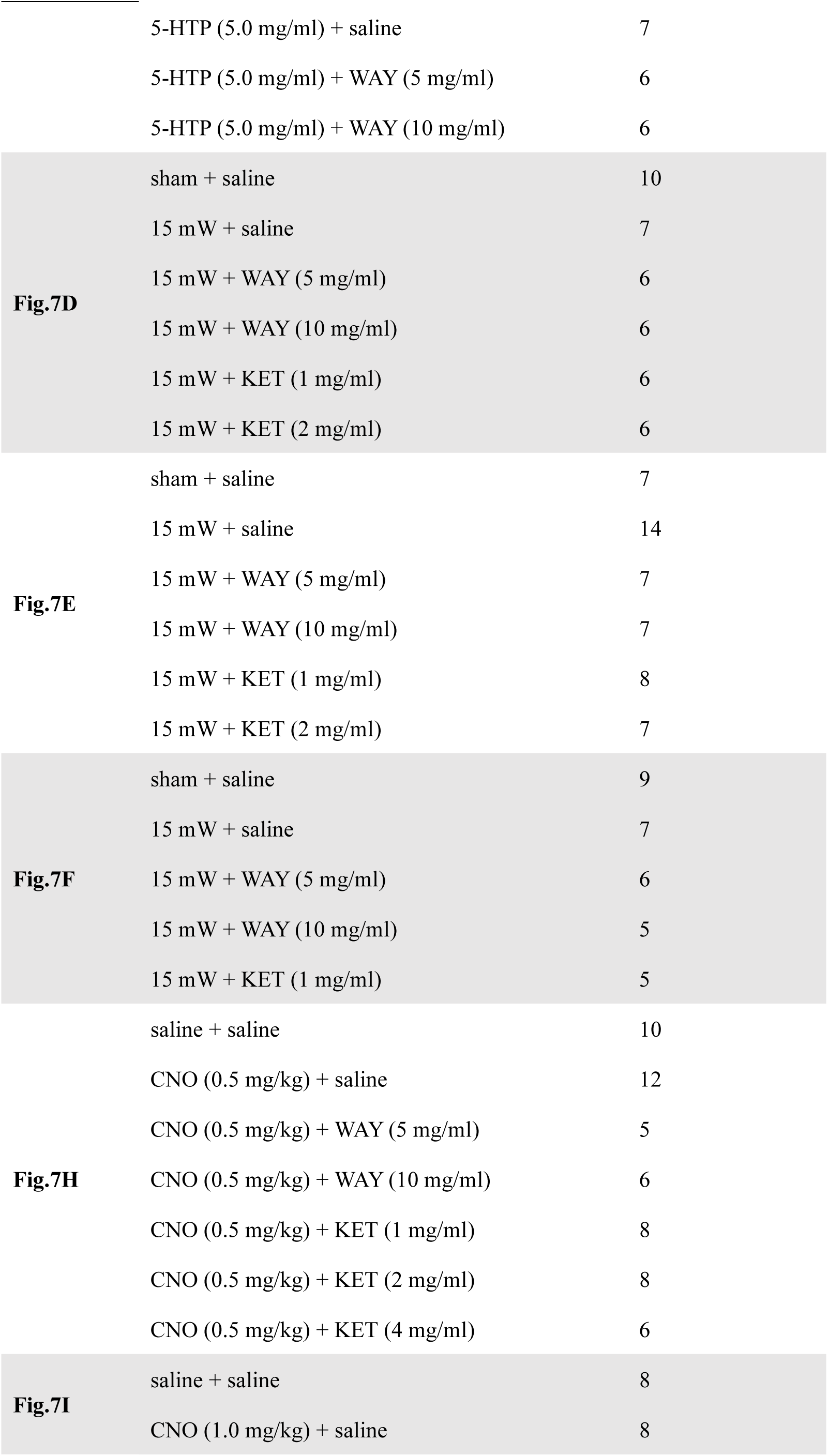
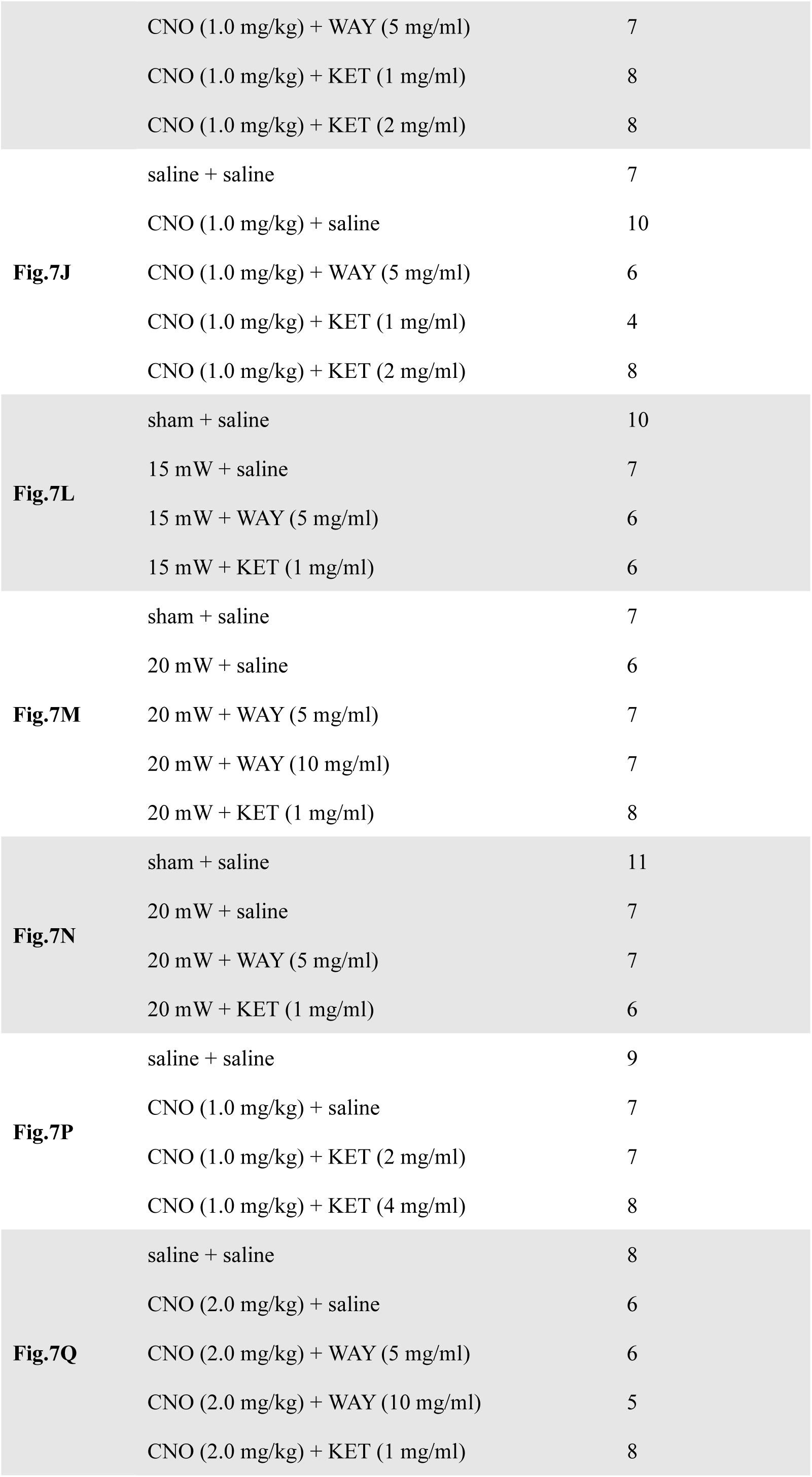
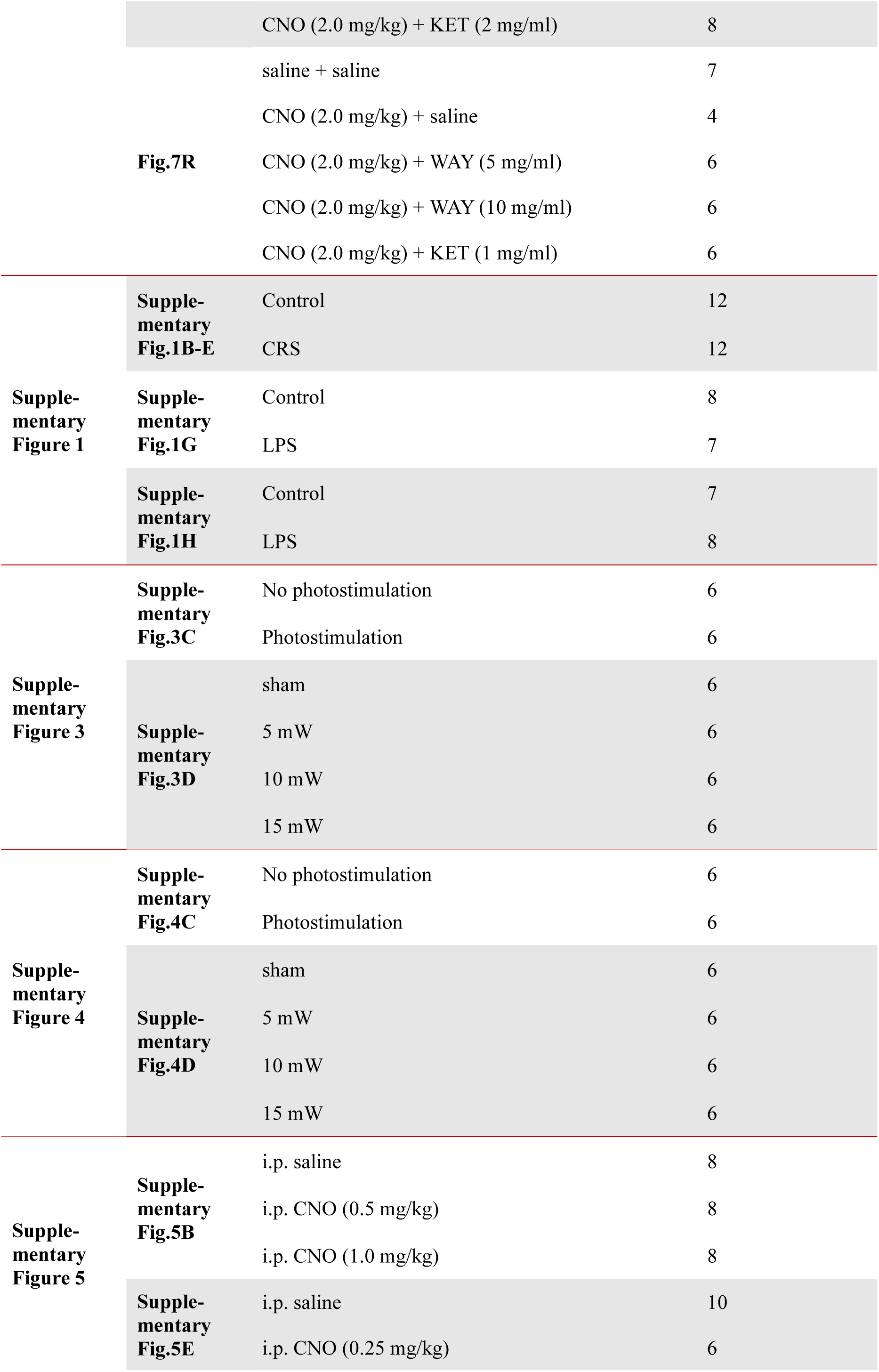
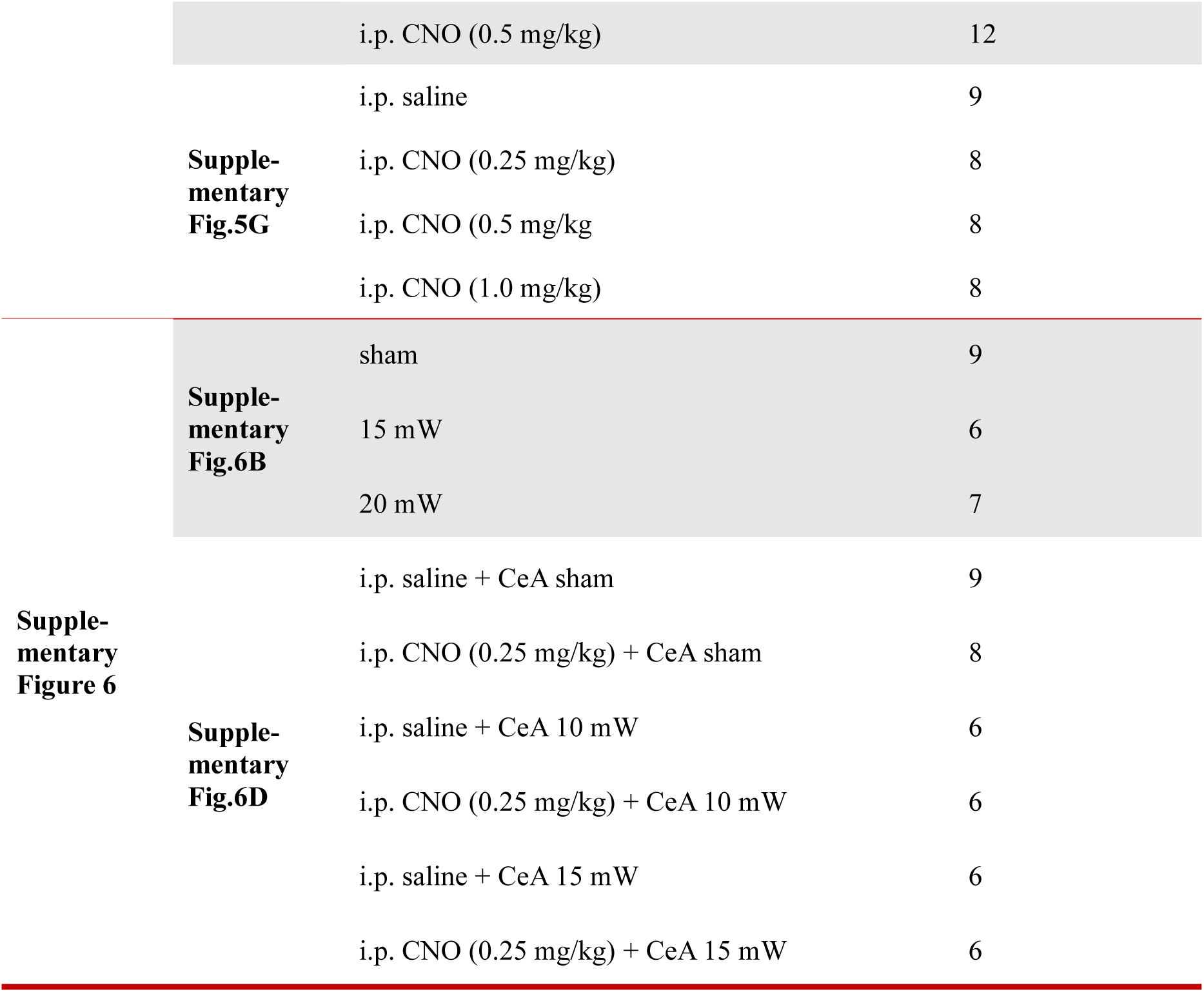
Summary of experimental groups of mice.

**Supplementary Table 2.** Summary of the total number of mice.

| Figure | Numbers of mice |
| --- | --- |
| Figure 1 | 188 |
| Figure 2 | 78 |
| Figure 3 | 240 |
| Figure 4 | 12 |
| Figure 5 | 103 |
| Figure 6 | 190 |
| Figure 7 | 474 |
| Supplementary Figure 1 | 54 |
| Supplementary Figure 3 | 36 |
| Supplementary Figure 4 | 36 |
| Supplementary Figure 5 | 85 |
| Supplementary Figure 6 | 63 |
| Total numbers of mice | 1559 |

**Supplementary Table 3.** Statistical analysis in each group.

| Figure | Comparisons | Test |
| --- | --- | --- |
| **Figure 1** | Emergence time (Fig.1B) | Ordinary one-way ANOVA |
|  | Emergence time (Fig.1D, E, F, H, L) | Ordinary one-way ANOVA |
| **Figure 2** | $\Delta F/F$ peak (Fig.2C, D, E, I, J) | Ordinary one-way ANOVA |
| | $\Delta F/F$ peak (Fig.2H) | Unpaired t-test |
| **Figure 3** | c-Fos(+)/TPH2(+)/section (Fig.3C) | Unpaired t-test |
|  | c-Fos(+)/TPH2(+)/section (Fig.3I) | Ordinary one-way ANOVA |
|  | Emergence time | Ordinary one-way ANOVA |
| **Figure 5** | $\Delta F/F$ peak (Fig.5C, D, E, K, L) | Ordinary one-way ANOVA |
|  | Emergence time | Ordinary one-way ANOVA |
| | $\Delta F/F$ peak (Fig.5J) | Unpaired t-test |
| **Figure 6** | c-Fos(+)/TPH2(+)/section | Ordinary one-way ANOVA |
|  | Emergence time | Ordinary one-way ANOVA |
| **Figure 7** | Emergence time | Ordinary one-way ANOVA |
| **Figure 8** | Correlation analysis | Correlation & Simple Linear Regression |
|  | Emergence time | Ordinary one-way ANOVA |
| **Supple-**<br>**mentary**<br>**Figure 1** | Center entries | Unpaired t-test |
|  | Total distance (m) | Unpaired t-test |
|  | Open arm entries | Unpaired t-test |
|  | Time in open arm (m) | Unpaired t-test |
|  | Immobility time (s) | Unpaired t-test |
|  | Time in center zone (s) | Unpaired t-test |
| **Supple-**<br>**mentary**<br>**Figure 3** | c-Fos(+)/TPH2(+)/section | Unpaired t-test |
|  | Emergence time | Ordinary one-way ANOVA |
| **Supple-**<br>**mentary**<br>**Figure 4** | c-Fos(+)/TPH2(+)/section<br>Emergence time | Unpaired t-test<br>Ordinary one-way ANOVA |
| **Supple-**<br>**mentary**<br>**Figure 5** | Emergence time | Ordinary one-way ANOVA |
| **Supple-**<br>**mentary**<br>**Figure 6** | Emergence time | Ordinary one-way ANOVA |

**Supplementary Table 4.**
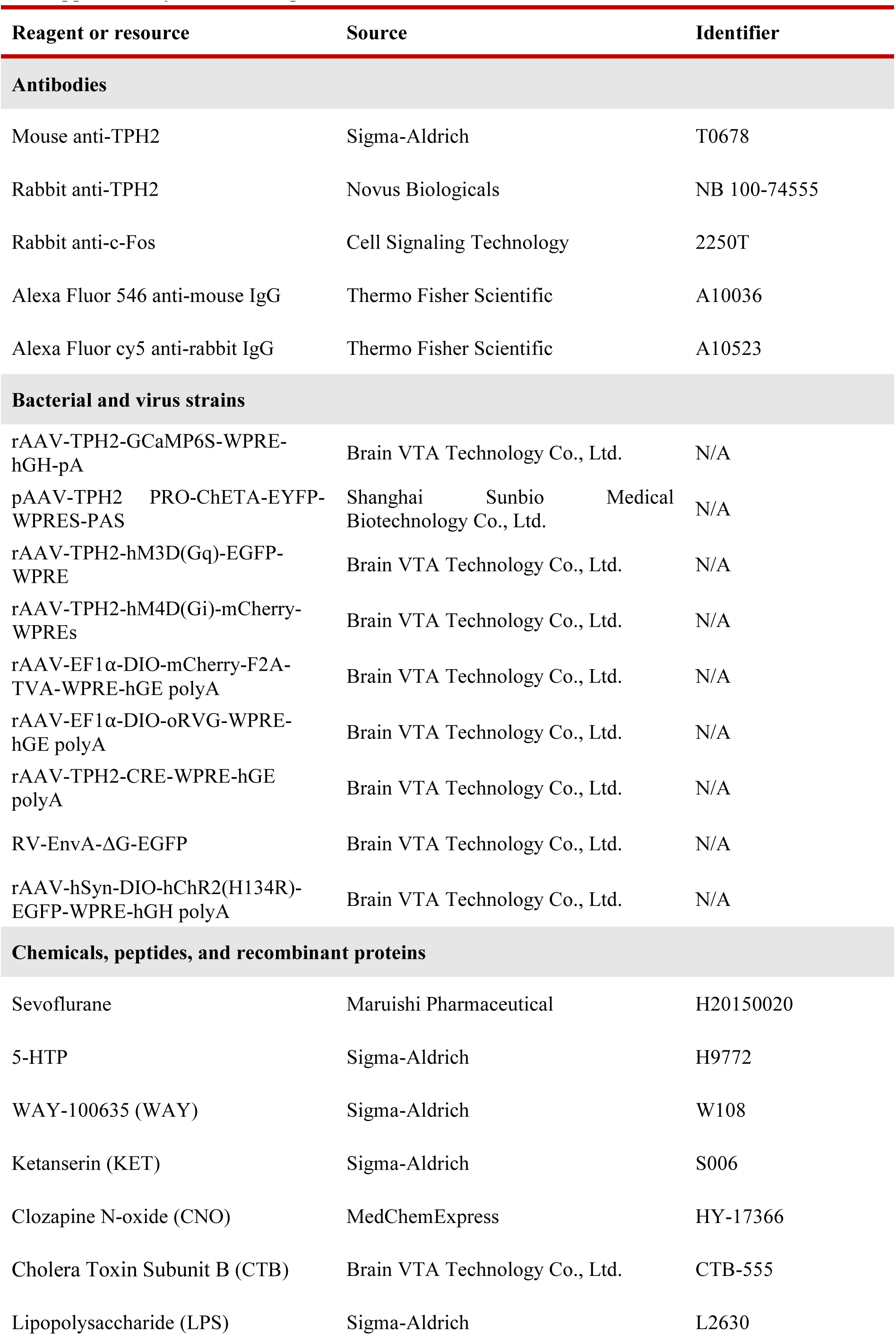
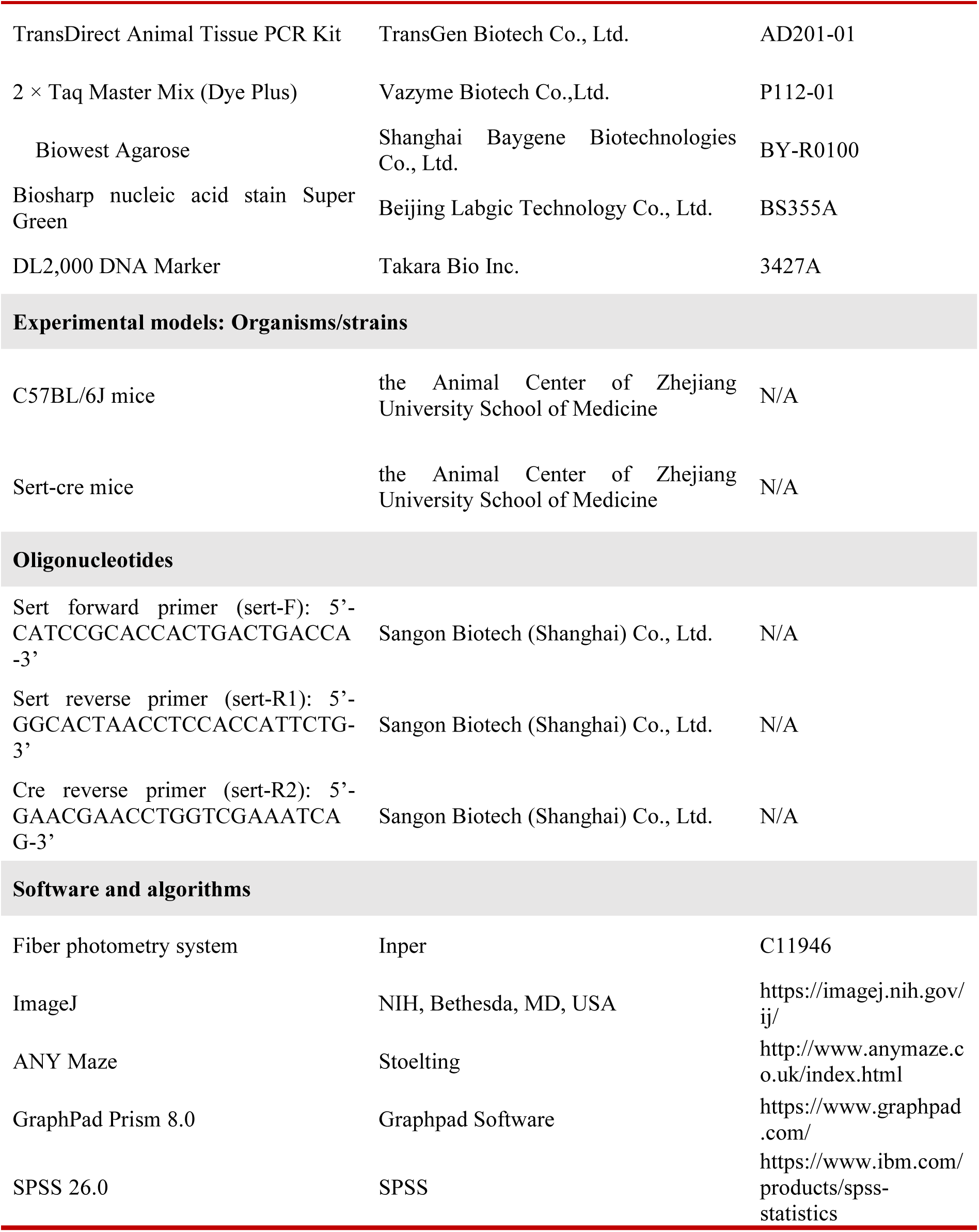
Reagent or resource.

